# Spindle-localized F-actin regulates polar MTOC organization and the fidelity of meiotic spindle formation

**DOI:** 10.1101/2025.05.07.652730

**Authors:** Edgar J. Soto-Moreno, Nourhan N. Ali, Florian Küller, Veselin Nasufovic, Michaela Frolikova, Olga Tepla, Jaromir Masata, Dirk Trauner, Amanda A. Patterson, Hans-Dieter Arndt, Katerina Komrskova, Magdalena Zernicka-Goetz, David M. Glover, Ahmed Z. Balboula

## Abstract

Mammalian oocytes are notoriously prone to chromosome segregation errors leading to aneuploidy. The spindle provides the machinery for accurate chromosome segregation during cell division. Mammalian oocytes lack centrioles and, therefore, the meiotic spindle relies on the organization of numerous acentriolar microtubule organizing centers into two poles (polar MTOCs, pMTOCs). The traditional view is that, in mammalian oocytes, microtubules are the sole cytoskeletal component responsible for regulating pMTOC organization and spindle assembly. We identified a novel F-actin pool that surrounds pMTOCs, forming F-actin cage-like structure. We demonstrated that F-actin localization on the spindle depends on unconventional myosins X and VIIb. Selective disruption of spindle-localized F-actin, using myosin X/VIIb knockdown oocytes or photoswitchable Optojasp-1, perturbed pMTOC organization, leading to unfocused spindle poles and chromosome missegregation. Here, we unveil an important function of F-actin in regulating pMTOC organization, a critical process for ensuring the fidelity of meiotic spindle formation and proper chromosome segregation.

## Introduction

Oocyte meiosis is a crucial process that involves a single round of DNA replication followed by two successive meiotic divisions. Around the time of birth, mammalian fetal oocytes are arrested at the dictyate stage of prophase I and remain in this state until puberty. After puberty, the oocyte within the preovulatory follicle resumes meiosis I and produces a haploid egg capable of fertilization. Despite its importance, oocyte meiosis I is a process that is highly associated with chromosome missegregation, leading to aneuploidy, the major genetic cause of miscarriage and congenital abnormalities^1–7^. Thus, understanding the regulatory mechanisms of oocyte meiosis I is essential to comprehend why this process is error prone.

The cell relies on the spindle machinery to properly segregate its chromosomes. In somatic cells, spindle assembly depends mainly on a pair of centriole-containing centrosomes. However, in mammalian oocytes, including human, centrioles are lost during early fetal development, and therefore lack classic centrosomes^8, 9^. In human prophase I-arrested oocytes, a single microtubule organizing center (MTOC), termed as human oocyte MTOC (huoMTOC), is initially located at the oocyte cortex prior to its migration towards the nucleus before nuclear envelope breakdown (NEBD). Upon NEBD, the perinuclear huoMTOC is reported to fragment into numerous MTOCs that are recruited to kinetochores^10^ to initiate chromatin-dependent Ran-GTP-mediated MT nucleation and spindle assembly^11, 12^. Prophase I-arrested mouse oocytes contain perinuclear MTOCs (termed class I). Upon meiotic resumption, sequential involvement of polo-like kinase (PLK1), dynein and Eg5 kinesin mediate perinuclear MTOC fragmentation into numerous MTOCs^13^. After NEBD, these numerous perinuclear class I MTOCs together with a subset of cytoplasmic MTOCs (termed class II) are organized^13, 14^ to promote MT nucleation (regulated by Aurora Kinase A and PLK1^15^) and to form the spindle poles (polar MTOCs, pMTOCs). Another class of cytoplasmic MTOCs (termed class III) remains in the cytoplasm until metaphase (metaphase cytoplasmic MTOCs, mcMTOCs) and does not contribute to spindle formation, yet plays a critical role in regulating spindle positioning and timely spindle migration^16^. Following NEBD, pMTOCs (*i.e.* MTOCs located at the spindle and contributed to spindle pole formation) undergo two critical steps to assemble a bipolar spindle with focused spindle poles^17–19:^ pMTOC clustering (*i.e.* MTOC-MTOC aggregation) and pMTOC sorting (*i.e.* MTOC movement and distribution towards the spindle poles). However, the molecular mechanism regulating pMTOC clustering and sorting to form focused spindle poles is largely unknown.

Filamentous actin (F-actin) is a cytoskeletal component that participate in different functions in mammalian oocytes during meiosis I including nucleus positioning^20–22^, chromosome capture and clustering^23^, kinetochore fiber (K-fiber) stability^28^, spindle positioning and spindle migration^24–27^. It has been widely accepted that, in mammalian oocytes, MTs, along with their associated proteins, constitute the only cytoskeletal elements responsible for regulating spindle bipolarity^13, 18, 19^. Recent data indicate that F-actin regulates bipolar spindle formation in human oocytes^29^. How F-actin interacts with MTs to regulate spindle bipolarity is unknown. F-actin has 2 distinct localizations in mammalian oocytes: at the cortex and cytoplasm. In both human and mouse oocytes, cytoplasmic F-actin is recruited to the MT spindle, through yet an unknown mechanism^24, 25, 28, 29^. Similarly, several laboratories described the presence of actin filaments in mitotic spindles^30, 31^. To date, there is no efficient tool to disrupt F-actin selectively at the spindle, and therefore the biological significance of spindle-localized F-actin pool remains largely unknown.

Here, we knockdown (KD) myosin X (MyoX) to disperse spindle-associated actin and employ a photoswitchable jasplakinolide to interfere with F-actin depolymerization dynamics to demonstrate that F-actin participates in regulating pMTOC organization to ensure the fidelity of meiotic spindle formation.

## Results

### F-actin regulates pMTOC sorting and clustering

To investigate the relationship between F-actin and pMTOC organization during oocyte meiosis I, we generated and employed Cep192-eGFP reporter mice^16^, in which expression and function of endogenous Cep192, an integral MTOC component, are neither perturbed nor overexpressed. Thus, Cep192-eGFP oocytes provide a highly suitable model for investigating pMTOC behavior during oocyte meiosis, while preserving their natural kinetics. To track pMTOC behavior in relation to F-actin, living Cep192-eGFP reporter oocytes were imaged using high-resolution time-lapse confocal microscopy in the presence of a low concentration of SiR-actin to visualize F-actin. At prometaphase I, we observed the presence of interconnected F-actin cables closely associated with pMTOCs (Fig. 1A and Supplementary Movie 1). Super-resolution Stimulated Emission Depletion (STED) microscopy confirmed the presence of this F-actin enrichment at pMTOCs in early metaphase I oocytes (Fig. 1B). Indeed, F-actin fluorescence intensity at pMTOCs (within 2 μm distance surrounding pMTOCs) was significantly higher than that at the spindle (Fig. 1C). To visualize how pMTOC-associated F-actin forms, we imaged both F-actin and pMTOCs for a relatively longer time during meiosis I. In full-grown prophase I-arrested (germinal vesicle, GV) oocytes, numerous perinuclear actin foci exist. After NEBD, we found that these perinuclear actin foci underwent a decrease in number and volume (Fig. 1D and Supplementary Fig. 1A,B). This decrease of actin foci coincided with a transient formation of an actin cloud at the spindle. At prometaphase I, in addition to previously reported actin fibers pulled from the cytoplasm to the spindle^28^, we observed the formation of de novo actin fibrils within the spindle area followed by their elongation (F-actin polymerization) to form F-actin fibers which start to associate with pMTOCs. (Supplementary Fig. 1C). Given that pMTOCs are associated with high MT density, actin fibril fluorescence intensity at pMTOCs was significantly higher than that at the spindle, leading ultimately to the formation of pMTOC-associated F-actin (Fig. 1D,E and Supplementary Fig. 1D).

**Figure 1:**
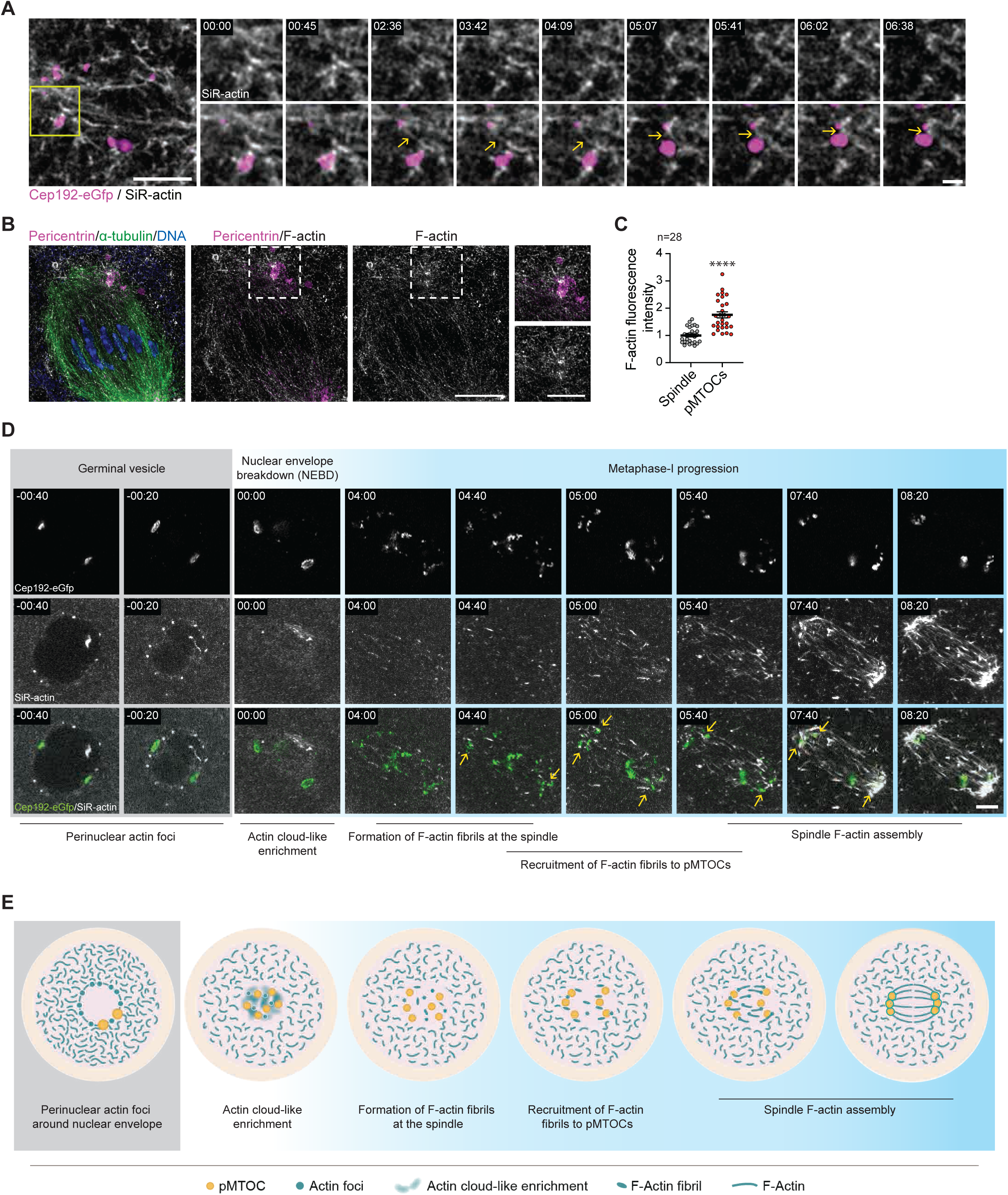
F-actin localizes to pMTOCs in mouse oocytes. (**A**) Time-lapse live confocal microscopy of pMTOCs (Cep192-eGfp, pseudo-magenta) and F-actin (SiR-actin, gray) in prometaphase I mouse oocytes. Fluorescence images were taken every 3 sec. Yellow arrows represent F-actin fibers connecting pMTOCs. Scale bar represents 5 μm (1.5 μm in zoomed images). Time points represent the timing from the start of confocal imaging. (**B**) Representative Super-resolution Stimulated Emission Depletion (STED) image of F-actin enrichment at pMTOCs in a metaphase I oocyte. The oocyte was labeled with phalloidin (F-actin, gray) α-tubulin antibody (green) and pricentrin antibody (magenta). DNA was stained with Hoechst (blue). Scale bar represents 10 μm. (**C**) Quantification of F-actin enrichment at pMTOCs. (**D**) Time-lapse live confocal microscopy of an oocyte expressing Cep192-eGfp (pMTOCs, pseudo-green) and stained with SiR-actin (F-actin, gray). Time points represent the timing from NEBD. Yellow arrows represent F-actin enrichment at pMTOCs. The total number of examined oocytes from 3 independent replicates is 24 oocytes. (**E**) Schematic diagram depicting the recruitment of F-actin to pMTOCs and the spindle, forming spindle-localized F-actin. Data are represented as means ± SEM. Asterisks represent significant differences, **** p<0.0001. The total number of analyzed oocytes per each experimental group (from 3 independent replicates) is specified above each graph.

Consistent with previous reports^13, 14, 17, 19^, we found that, after NEBD, pMTOCs undergo a transient fragmentation followed by two overlapping processes of pMTOC clustering (*i.e.* MTOC-MTOC aggregation) and pMTOC sorting (*i.e.* MTOC movement and distribution towards the poles) to assemble the spindle poles (Fig. 2A-C). Accordingly, upon pMTOC clustering, average pMTOC numbers decreased gradually from ∼26 pMTOCs post-NEBD to ∼9 pMTOCs at metaphase I (Fig. 2B), whereas the average pMTOC volume increased from 15.8 ± 1.79 μm^3^ to 31.5 ± 1.36 μm^3^ at metaphase I (Fig. 2C). To perturb F-actin, we treated oocytes with cytochalasin D (CytoD) at NEBD and, similar to control oocytes, we observed a transient increase in numbers and decrease in volumes of pMTOCs at ∼1 h post-NEBD (Fig. 2B,C), indicating that F-actin does not play a role in regulating post-NEBD pMTOC fragmentation. Strikingly, compared to control oocytes, inhibition of F-actin at NEBD impaired proper pMTOC clustering and sorting towards the spindle poles (Fig. 2A-C). To characterize this response in detail, we conducted immunocytochemistry to label MTOCs and the spindle using antibodies against pericentrin (an integral component of MTOCs in mouse oocytes^32^) and α-tubulin, respectively. Inhibition of F-actin polymerization by CytoD or latrunculin B (LatB) treatment at NEBD resulted in pMTOC clustering defects [as evidenced by an increase in pMTOC numbers (Fig. 2D,G) and a decrease in pMTOC volumes (Fig. 2D,H)] and pMTOC sorting defects [as evidenced by the inappropriate scattering of pMTOCs along the spindle axis (Fig. 2D,E) and the decreased ratio of the total volume of pMTOCs at the spindle poles to the total volume of all pMTOCs in the spindle (Fig. 2D,F)] at metaphase I (5 h post-NEBD) when compared to DMSO-treated oocytes. This pMTOC clustering defects may explain the previous observation of reduced fluorescence intensity of pMTOCs in CytoD-treated oocytes (likely due to decreased pMTOC volume below the imaging detection threshold)^33^. It is unlikely that the pMTOC clustering and sorting defects in F-actin-perturbed oocytes are due to a cell cycle delay because (1) time-lapse confocal microscopy revealed that F-actin-perturbed oocytes attempt to divide symmetrically with unclustered and unsorted pMTOCs (Fig. 2A), and (2) arresting oocytes at metaphase I for a longer period (8 h post-NEBD, the timing at which the oocytes typically progress beyond metaphase I) using MG-132, a cell-permeable proteosome inhibitor, did not prevent pMTOC sorting and clustering defects in F-actin-perturbed oocytes (Supplementary Fig. 1E-I). To investigate whether dynamic F-actin is required for regulating pMTOC sorting and clustering, we treated oocytes (at NEBD) with jasplakinolide (Jasp, a cell permeable F-actin stabilizing agent). Strikingly, stalling F-actin dynamics by using Jasp phenocopied the pMTOC sorting (Supplementary Fig. 1J-L) and clustering (Supplementary Fig. 1J,M,N) defects observed in CytoD- and LatB-treated oocytes, indicating that dynamic behavior of F-actin fibers is required for pMTOC sorting and clustering in oocytes. Although we did not observe defective spindle bipolarity (Fig. 2D) or an altered spindle length to width ratio (Supplementary Fig. 2A), we found a significant increase of spindle pole width in F-actin-perturbed oocytes when compared to controls (Fig. 2I), indicative of unfocused spindle poles (Fig. 2D), a phenomenon that correlates with chromosome missegregation^34–41^. Moreover, consistent with the finding that F-actin perturbation had no effect on MT dynamics in metaphase II mouse oocytes^28^, we did not observe altered pMTOC-associated MT dynamics in F-actin-perturbed oocytes as assessed by Fluorescence Recovery After Photobleaching (FRAP assay, Supplementary Fig. 2B,C). Furthermore, in contrast to control oocytes, we frequently observed a higher incidence of pMTOC-associated MTs located outside the dominant spindle pole area (*i.e.* greater than 5 μm distance from the dominant spindle pole center) in F-actin-perturbed oocytes (Fig. 2D,J). Thus, our results indicate that F-actin regulates pMTOC sorting and clustering in mouse oocytes to ensure the fidelity of meiotic spindle construction. Because the spindle provides the machinery for faithful chromosome segregation, we examined chromosome segregation in F-actin-perturbed oocytes. We found a significant increase in the incidence of chromosome misalignment at metaphase I and lagging chromosomes at anaphase/telophase I in CytoD-, LatB- and Jasp-treated oocytes when compared to control oocytes (Supplementary Fig. 2D-G). A previous study showed that disruption of F-actin by CytoD leads to chromosome missegregation by decreasing K-fiber stability at late metaphase I^28^. To determine whether the effect of CytoD on pMTOC clustering and sorting leads to chromosome missegregation independently of its role in K-fiber stability, we examined K-fiber stability and pMTOC clustering and sorting after CytoD washout (i.e. F-actin is disrupted only at 1-4 h post-NEBD, the period at which the majority of pMTOCs are clustered and sorted). We found that acute disruption of F-actin resulted in pMTOC clustering and sorting defects at late metaphase I (Supplementary Fig 3A-D) and increased lagging chromosomes at anaphase/telophase I (Supplementary Fig 3E,F) even in the presence of normal K-fiber stability (Supplementary Fig 3G,H). These results suggest that pMTOC clustering and sorting defects induced by F-actin disruption can lead to chromosome segregation errors independent of F-actin role on K-fiber stability.

**Figure 2:**
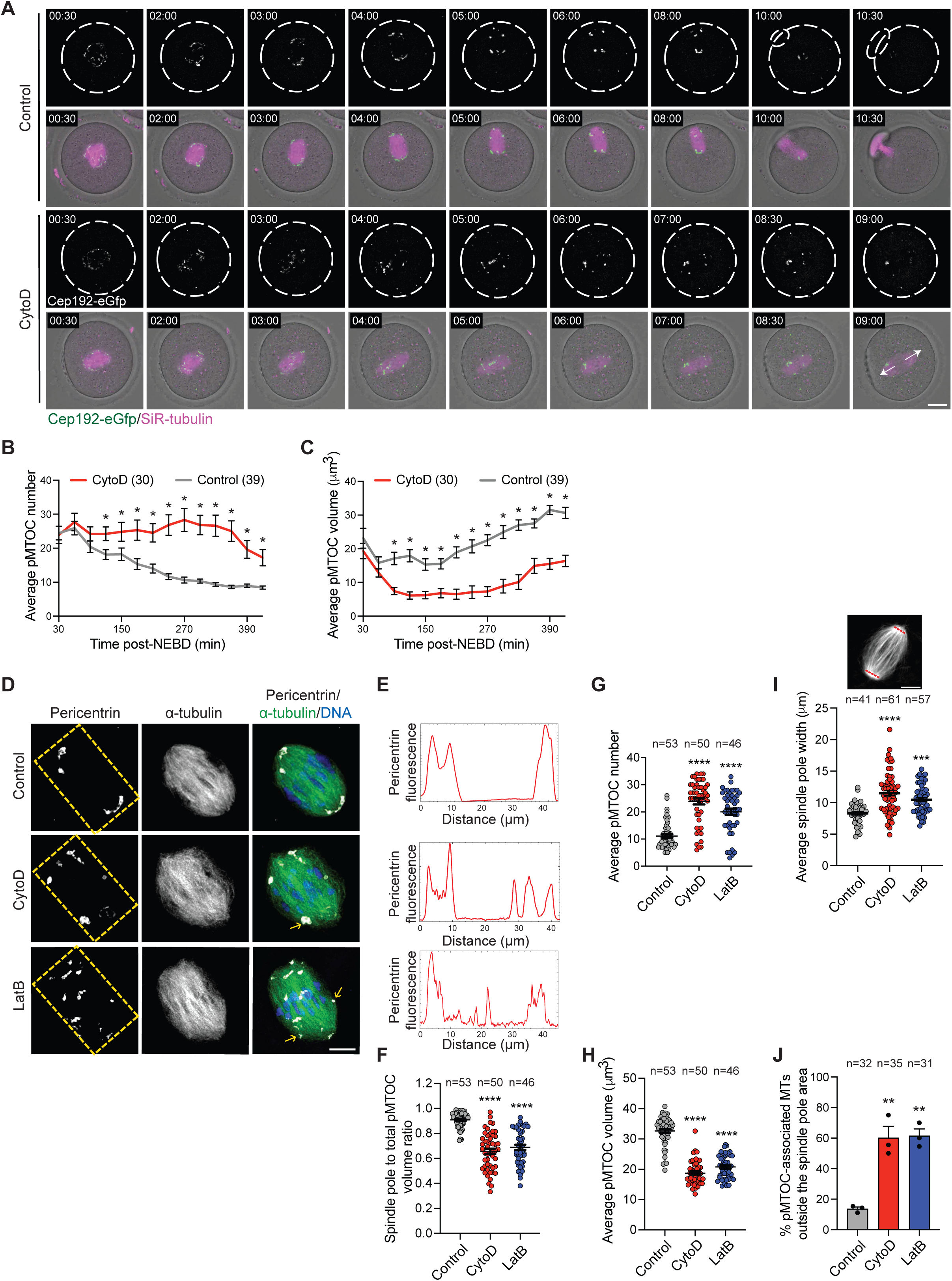
F-actin regulates pMTOC sorting and clustering in mouse oocytes. (**A**) Time-lapse live confocal microscopy of pMTOCs (Cep192-eGfp, green) and microtubules (SiR-tubulin, magenta) in mouse oocytes treated with DMSO (control) or Cytochalasin D (CytoD) after NEBD. Fluorescence images were taken every 30 min. Scale bar represents 20 μm. (**B**) Average pMTOC number quantification after 3D reconstruction from oocytes in **A.** (**C**) Average pMTOC volume quantification after 3D reconstruction of Cep192-eGfp from oocytes in **A**. (**D**) Representative confocal images of metaphase I oocytes treated with DMSO (control), CytoD, or latrunculin B (LatB) at NEBD. Metaphase I oocytes were immunostained with α-tubulin to label the spindle (green) and pericentrin to label MTOCs (gray). DNA was labeled with 4′,6-diamidino-2-phenylindole, dihydrochloride (DAPI, blue). Yellow arrows represent pMTOC-associated microtubules outside the spindle pole area. Scale bar represents 10 μm. (**E**) Plot profile of pMTOCs (pericentrin fluorescence) along the longitudinal axis of the spindle as represented by the yellow rectangle area in **D**. (**F**) Quantification of pMTOC volume at the spindle poles to the total volume of all pMTOCs in the spindle. (**G**) Average pMTOC number quantification in **D**. (**H**) Average pMTOC volume quantification in **D**. (**I**) Average spindle pole width of oocytes treated with DMSO (control), CytoD, or LatB. The spindle pole positions were determined as the endpoints of the spindle. A line was drawn that is passed through and spans the dominant spindle pole(s) at each side and averaged to calculate the average spindle pole width for each oocyte. (**J**) Average percentage of oocytes with pMTOC-associated microtubules outside the spindle pole area. The numbers and volumes of pMTOCs were auto calculated following the 3D reconstruction of pMTOCs. Data are represented as means ± SEM. Asterisks represent significant differences, * p<0.05, ** p<0.01, *** p<0.001, and **** p<0.0001. The total number of analyzed oocytes per each experimental group (from 3 independent replicates) is specified above each graph.

These phenotypes of F-actin-perturbed mouse oocytes accord with the disrupted spindle morphology^29^, altered spindle pole focusing and NuMA (nuclear mitotic apparatus protein that is associated with MT minus-ends) clustering and sorting defects seen when F-actin is perturbed in bovine (Supplementary Fig. 4A,B), porcine (Supplementary Fig. 4C,D) and human oocytes (Supplementary Fig. 4E,F). It is of note that, in human oocytes, NuMA foci are clustered at the spindle poles and their depletion resulted in unfocused poles^12^, whereas in mouse oocytes where spindle pole focusing is governed by pMTOCs, NuMA depletion resulted in an opposite phenotype-hyperfocused spindle poles^40^.

### Spindle-localized F-actin regulates pMTOC sorting and clustering

Following NEBD, F-actin has two distinct localizations in mammalian oocytes: at the cortex and in the cytoplasm. Cortical F-actin nucleation is regulated by the ARP2/3 complex which localizes at the oocyte cortex^42, 43^, whereas cytoplasmic F-actin nucleation is primarily depends on Formin 2 (Fmn2) and SPIRE 1/2^44–46^. As previously reported, Fmn2 localizes at the cytoplasm and enriched at the cortex and the spindle (Supplementary Fig. 5A). To investigate whether the cortical or cytoplasmic F-actin pool regulates pMTOC clustering and sorting in mouse oocytes, we inhibited either Arp2/3 using CK-666 (a selective Arp2/3 inhibitor)^43, 47^ or Fmn2 using SMIFH2 (a small molecular inhibitor of the formin homology 2 domains^48^). The CK-666 treatment did not affect cytoplasmic F-actin (Supplementary Fig. 5B) and had no significant effect on pMTOC sorting (Supplementary Fig. 5C-E) and clustering (Supplementary Fig. 5C,G,G). In contrast, SMIFH2 treatment decreased cytoplasmic F-actin (Supplementary Fig. 5H) and phenocopied pMTOC sorting (Supplementary Fig. 5I-K) and clustering (Supplementary Fig. 5I,L,M) defects observed in F-actin-perturbed oocytes. In addition to Fmn2, SMIFH2 can inhibit other myosins including myosin II. We found that inhibition of myosin II by blebbistatin had no significant effect on cytoplasmic F-actin (Supplementary Fig. 5N) or pMTOC organization (Supplementary Fig. 5O-S), suggesting that SMIFH2 effect on spindle-localized F-actin and pMTOC organization is not due to myosin II inhibition. These findings reveal a novel role of F-actin in regulating two critical processes —pMTOC clustering and sorting— during oocyte meiosis, and suggest that the cytoplasmic F-actin pool interacts, yet through unknown mechanisms, with the MT spindle to carry out these functions. Although SMIFH2 (Supplementary Fig. 5H), CytoD and latB (Supplementary Fig. 5T-V) significantly decreased spindle-localized F-actin and Jasp perturbed F-actin dynamics at the spindle without affecting G-actin level (Supplementary Fig. 5W-Z), their effect is global throughout the entire oocyte. We therefore wished to ask whether the spindle-localized F-actin pool regulates pMTOC sorting and clustering and so sought an experimental tool to selectively perturb F-actin function at the spindle. F-actin localization to the spindle is MT dependent^28^. The unconventional MyoX is an F-actin-based motor protein with a myosin tail homology 4 (MyTH4) domain in its C-terminal region, which enables it to bind MTs directly, giving the potential to link F-actin and MTs^49, 50^. Moreover, MyoX localizes to the spindle pole in mitotic cells and to the spindle MTs in *Xenopus* eggs, and, in both cases, its inhibition results in spindle defects^51–54^ likely through perturbing the link between spindle-intrinsic forces and actin-dependent cortical forces. Using Western blot analysis, we confirmed that MyoX is expressed in mouse oocytes (Supplementary Fig. 6A). MyoX localized to the meiotic spindle and enriched at the pMTOCs at prometaphase I (Supplementary Fig. 6B) and metaphase I (Supplementary Fig. 6C), as previously reported^55, 56^.

Treating oocytes with nocodazole, a MT depolymerizing drug, decreased MyoX’s localization to the spindle and pMTOCs (Supplementary Fig. 6C), indicating a dependence on MTs for the localization of MyoX to the meiotic spindle and pMTOCs. To determine whether MyoX recruits F-actin to the meiotic spindle, we sought to deplete MyoX by siRNA-mediated KD. The microinjection of full-grown GV oocytes with a cocktail of siRNAs, targeted to different regions of the MyoX gene, significantly decreased *MyoX* mRNA transcript abundance (Supplementary Fig. 6D) and the level of MyoX at metaphase I (Supplementary Fig. 6E,F). In contrast to control oocytes, in which F-actin localizes to the spindle, MyoX KD significantly decreased spindle-localized F-actin (Fig. 3A,B) without altering cytoplasmic or cortical F-actin levels (Fig. 3A,C,D). In addition, MyoX KD significantly disrupted the alignment of F-actin fibers along the longitudinal axis of the spindle, compared to controls (Fig. 3A,E). These findings suggest that MyoX plays an important role in recruiting F-actin to the spindle and that MyoX KD represents an efficient tool to selectively perturb spindle-localized F-actin. Importantly, siRNA-mediated KD of MyoX phenocopied the pMTOC sorting and clustering defects in F-actin-perturbed oocytes (Fig. 3F-J), suggesting that spindle-localized F-actin plays a role in regulating pMTOC organization during oocyte meiosis. The microinjection of siRNA-resistant MyoX cRNA rescued both pMTOC sorting and clustering defects (Supplementary Fig. 6G-K) and the reduced spindle-localized F-actin (Supplementary Fig. 6G,L) in MyoX KD oocytes, confirming the specificity of MyoX siRNA. When MyoX KD oocytes were allowed to mature, we did not observe defective spindle migration to the oocyte cortex or cytokinesis failure relative to controls (Fig 3J and Supplementary Fig. 7A,B), further confirming that cytoplasmic and cortical F-actin were not perturbed in MyoX KD oocytes. Consistent with several reports indicating that unfocused spindle poles and/or unclustered pMTOCs result in chromosome segregation defects^34–41, 57^, we observed a significant increase in the incidence of chromosome misalignment and lagging chromosomes in MyoX KD oocytes when compared to control oocytes (Fig. 3K,L,N). To confirm the role of MyoX in F-actin recruitment to the spindle and pMTOC organization, we wished to perturb MyoX and spindle-localized F-actin using a different approach. To this end, we generated a dominant negative mouse MyoX mutant that lacks MT binding ability in its MyTH4 domain (MyoX MyTH4-mutant, K1651D/K1654D)^58^. Expectedly, full-length MyoX localized to the spindle including pMTOCs (Supplementary Fig. 7C) and increased spindle-localized F-actin (Supplementary Fig. 7E,F), indicating that MyoX promotes F-actin localization at the spindle. In contrast to full-length MyoX, MyoX MyTH4-mutant failed to localize properly to either the spindle or pMTOCs (Supplementary Fig. 7C) and resulted in significantly decreased F-actin localization to the spindle (Supplementary Fig. 7C,D). Time-lapse microscopy confirmed that the expression of the MyoX MyTH4 dominant-negative mutant or MT depolymerization by nocodazole significantly decreased F-actin localization at the spindle and pMTOCs (Supplementary Fig. 7G-I). Since MyoX carries out its function while being in MyoX dimers^59^, it is plausible that the MyoX MyTH4 mutant exerts a dominant negative effect by creating non-functional dimers with endogenous MyoX. This likely explains why spindle-localized F-actin was not completely abolished in MyoX MyTH4-mutant-expressing oocytes. Together, these observations suggest that MyoX localization to the spindle depends on its MyTH4 domain for MT interaction and that such MyoX localization is essential for F-actin recruitment to the spindle. Importantly, consistent with siRNA-mediated MyoX KD, expression of the MyoX MyTH4 dominant-negative mutant resulted in pMTOC sorting (Fig. 3O-Q) and clustering (Fig. 3O,R,S) defects. Thus, MyoX regulates pMTOC sorting and clustering in mouse oocytes. To confirm whether reduced spindle-localized F-actin is the cause of pMTOC sorting and clustering defects in MyoX perturbed oocytes, we next investigated whether forcing F-actin localization to the spindle in oocytes expressing the MyoX MyTH4-mutant could rescue, at least in part, the observed defects in pMTOC sorting and clustering. To this end, we expressed microtubule-associated protein 4 (MAP4) fused to the calponin homology domain of human Utrophin (Utr-CH, which binds F-actin), a construct which has been shown to force F-actin localization to the spindle^28^. It is of note that Utr-CH binds F-actin without stabilizing it^60^. For this reason, it is commonly used as a reliable tool to study F-actin distribution and dynamics in different living cell systems including mammalian oocytes^23–25, 61^. Accordingly, the expression of Utr-CH-MAP4 enriched F-actin on the spindle even in the absence of MyoX (Supplementary Fig. 8A). Importantly, forcing F-actin localization to the spindle using a low concentration of Utr-CH-MAP4, which does not perturb F-actin dynamics compared to a high concentration of Utr-CH-MAP4 (Supplementary Fig. 8B,C), significantly rescued the clustering and sorting defects observed in MyoX KD oocytes (Supplementary Fig. 8D-H) and MyoX MyTH4-expressing oocytes (Fig. 3O-S). These results strongly suggest that MyoX acts upstream of F-actin recruitment and that the spindle-localized F-actin pool is indispensable for efficient MTOC sorting and clustering.

**Figure 3:**
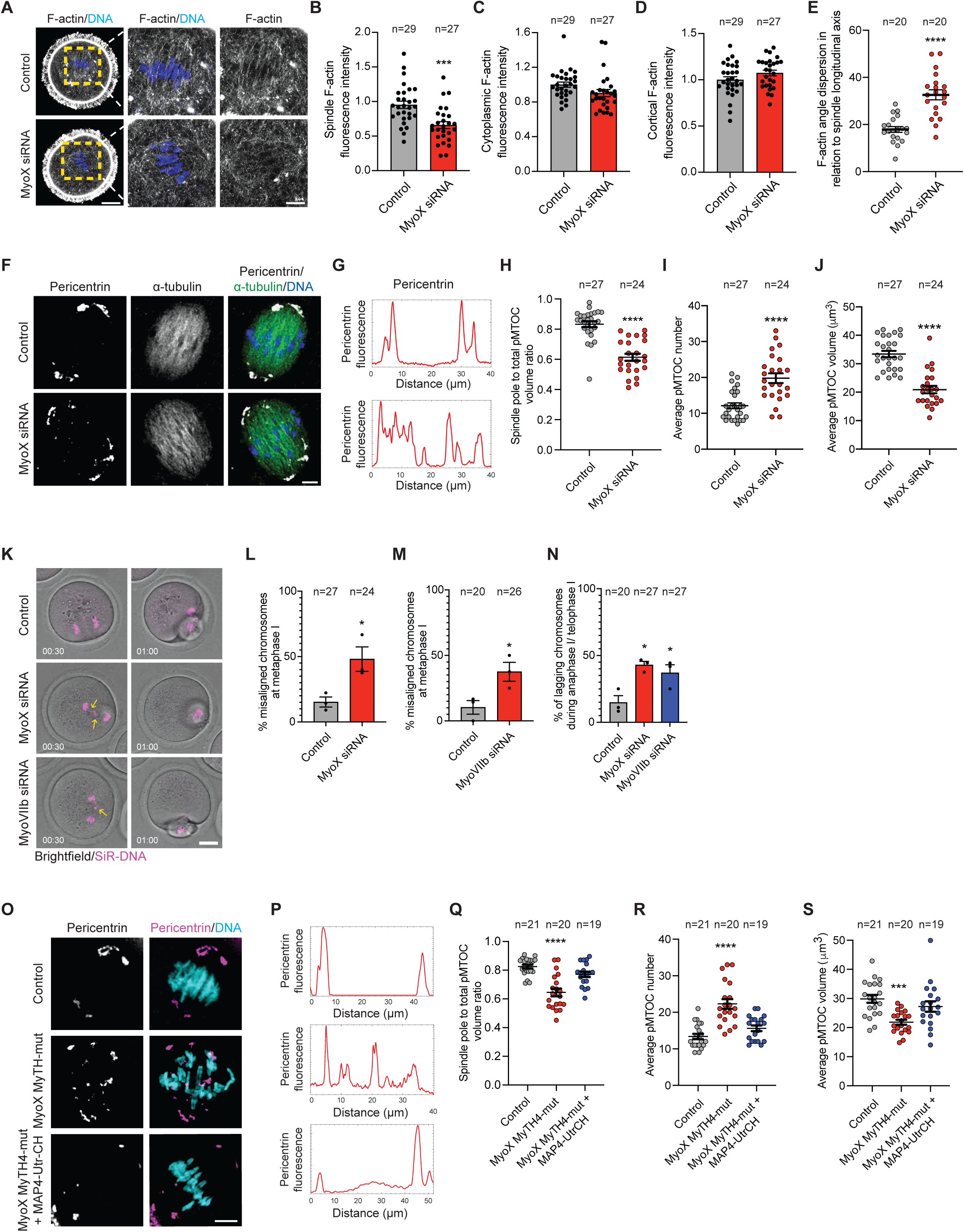
Myosin X regulates spindle-localized F-actin and pMTOC organization in mouse oocytes. (**A,F**) Representative confocal images of metaphase I oocytes microinjected with control siRNA or Myosin X siRNA at the germinal vesicle stage. The oocytes were stained with phalloidin (F-actin) and 4′,6-diamidino-2-phenylindole, dihydrochloride (DAPI) in **A** or immunostained with α-tubulin and pericentrin antibodies in **F.** Scale bar represents 10 μm. (**B**) Quantification of average spindle-localized F-actin. (**C**) Quantification of average cytoplasmic F-actin. (**D**) Quantification of average cortical F-actin. (**E**) Quantification of F-actin angle dispersion in relation to the longitudinal axis of the spindle. (**G**) Plot profile of pMTOCs (pericentrin fluorescence) along the longitudinal axis of the spindle. (**H**) Quantification of pMTOC volume at the spindle poles to the total volume of all pMTOCs in the spindle. (**I**) Average pMTOC number quantification. (**J**) Average pMTOC volume quantification. (**K)** Metaphase I oocytes microinjected with control siRNA, Myosin X siRNA or Myosin VIIb siRNA were stained with SiR-DNA followed by time-lapse confocal microscopy. Scale bar represents 20 μm. Chromosomes were tracked throughout live imaging by using the autotracking function of Imaris software. Yellow arrow heads represent lagging chromosomes. Time points represent the timing from the start of confocal imaging. (**L**) Quantification of average percentage of misaligned chromosomes at metaphase I. (**M**) Quantification of average percentage of misaligned chromosomes at metaphase I. (**N**) Average percentage of lagging chromosomes of control, Myosin X knockdown and Myosin VIIb knockdown oocytes. (**O**) Representative confocal images of metaphase I oocytes expressing mRuby (control), mRuby-Myosin X microtubule binding domain mutant (MyoX MyTH-mut) or MyoX MyTH-mut + MAP4-UtrCH-GFP cRNA. Metaphase I oocytes were immunostained with pericentrin to label MTOCs and stained with DAPI to label DNA. Scale bar represents 10 μm. (**P**) Plot profile of pMTOCs (pericentrin fluorescence) along the longitudinal axis of the spindle. (**Q**) Quantification of pMTOC volume at the spindle poles to the total volume of all pMTOCs in the spindle. (**R**) Average pMTOC number quantification after 3D reconstruction of pericentrin. (**S**) Average pMTOC volume quantification. Data are represented as means ± SEM. Asterisks represent significant differences, * p<0.05, *** p<0.001, and **** p<0.0001. The total number of analyzed oocytes (from 3 independent replicates) is specified above each graph.

Despite several in vitro studies indicating that MyoX is an indispensable protein for meiotic and mitotic divisions^51, 52, 62^, MyoX knockout (KO) mice appear to be fertile and to have normal mitosis^63^. Indeed, oocytes from MyoX KO mice have apparently normal spindle morphogenesis and positioning^64^, suggesting that MyoX function(s) at the spindle may be compensated by other unconventional myosins. Myosins VIIb and XVa are also unconventional actin-based motor proteins that contain MyTH4 and FERM domains that can bind to both actin and MTs ^65–69^. We hypothesized that MyoVIIb and/or MyoXVa may also cross-link F-actin and MTs. We first confirmed that MyoVIIb is expressed in mouse oocytes by Western blot analysis (Supplementary Fig. 6A). To investigate whether MyoVIIb and/or MyoXVa can regulate F-actin localization to the meiotic spindle, we knocked down MyoVIIb or MyoXVa and assessed F-actin localization to the spindle. siRNA-mediated KD of MyoVIIb or MyoXVa decreased the mRNA transcript abundance (Supplementary Fig. 9A,B) and protein levels of MyoVIIb and MyoXVa (Supplementary Fig. 9C-F), respectively. Interestingly, KD of MyoVIIb, but not MyoXVa, significantly decreased spindle-localized F-actin (Supplementary Fig. 9C,E,G) when compared to controls, indicating that MyoVIIb also regulates the recruitment of spindle-localized F-actin. We did not observe alteration in cytoplasmic F-actin (Supplementary Fig. 9H), spindle migration (Fig. 3K and Supplementary Fig. 7A) or cytokinesis (Fig. 3K and Supplementary Fig. 7B) in MyoVIIb or MyoXVa KD oocytes when compared to controls. Importantly, oocytes with reduced levels of MyoVIIb, but not MyoXVa exhibited perturbed pMTOC sorting (Supplementary Fig. 9I-K) and clustering (Supplementary Fig. 9I,L,M), suggesting that MyoVIIb could also regulate pMTOC organization and may explain the apparently normal spindle morphogenesis in MyoX KO oocytes^64^. The microinjection of siRNA-resistant MyoVIIb cRNA rescued both reduced spindle-localized F-actin and the pMTOC sorting and clustering defects in MyoVIIb KD oocytes, confirming the specificity of MyoVIIb siRNA (Supplementary Fig. 6G-L). We also observed a significant increase in the incidence of chromosome misalignment (Fig. 3M) and lagging chromosomes (Fig. 3K,N) in MyoVIIb KD oocytes when compared to control oocytes. We then investigated whether MyoX and MyoVIIb have overlapping functions in regulating spindle-localized F-actin. We found that MyoX and MyoVIIb double KD resulted in further depletion of spindle-localized F-actin when compared to MyoX or MyoVIIb single KD (Fig. 4A,B). We then asked whether MyoVIIb can rescue reduced spindle-localized F-actin and improper pMTOC sorting and clustering in MyoX KD oocytes. We found that full-length MyoVIIb localizes to the spindle and is enriched at pMTOCs, similar to MyoX localization (Fig. 4C). Importantly, overexpressing MyoVIIb rescued reduced F-actin localization to the spindle without affecting cytoplasmic F-actin (Fig. 4D,I,J) and the pMTOC sorting (Fig. 4D-F) and clustering (Fig. 4D,G,H) defects in MyoX KD oocytes. Taken together, our findings suggest that unconventional MyoX and MyoVIIb are required to ensure F-actin localization to the spindle and proper pMTOC organization during oocyte meiosis.

**Figure 4:**
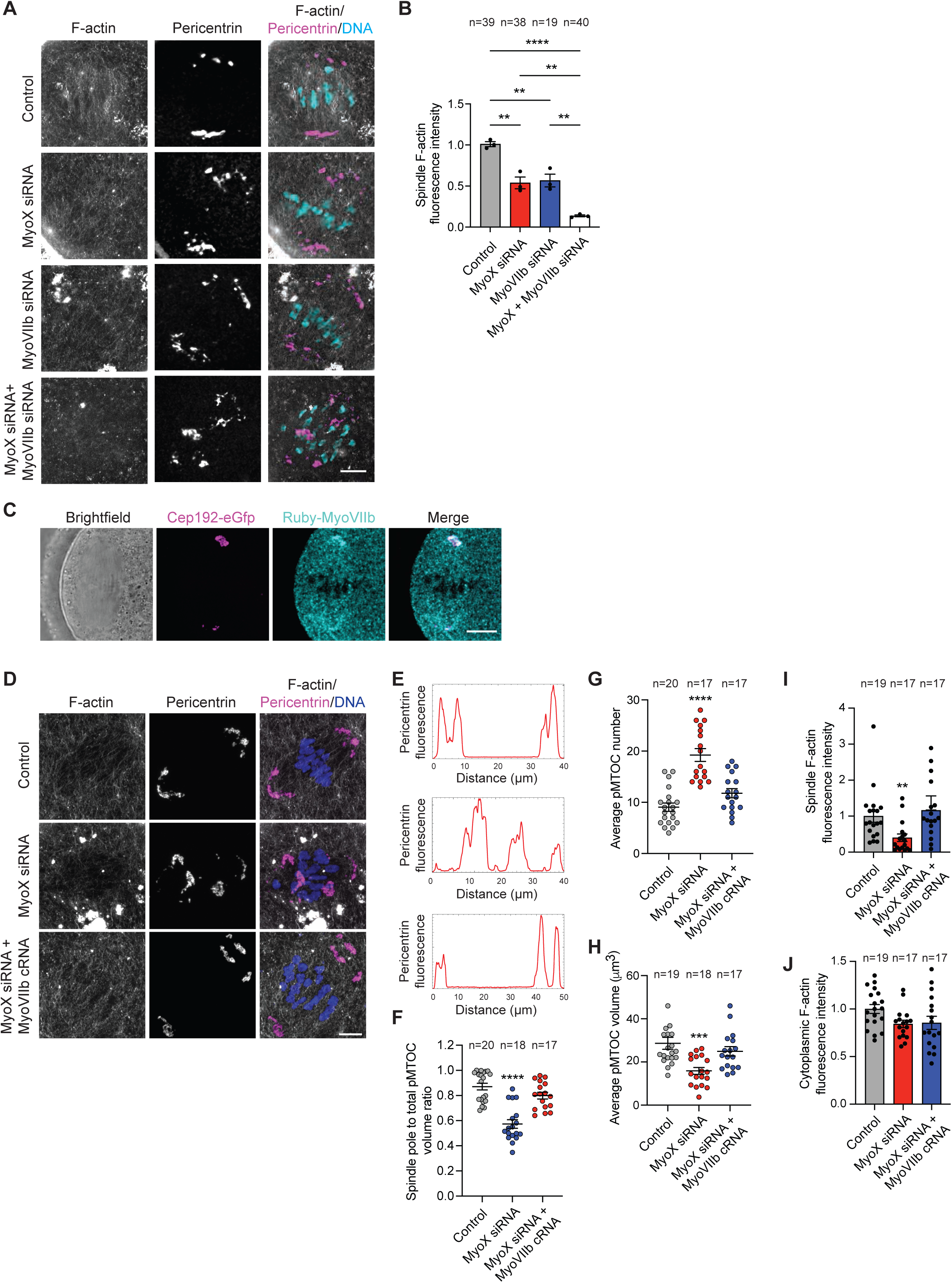
Myosin X and Myosin VIIb have overlapping functions in regulating spindle-localized F-actin and pMTOC organization. (**A**) Representative confocal images of metaphase I oocytes microinjected with control siRNA, Myosin X siRNA, Myosin VIIb siRNA or Myosin X siRNA + Myosin VIIb siRNA at the germinal vesicle stage. The oocytes were immunostained with pericentrin antibody (MTOCs) and stained with phalloidin (F-actin) and 4′,6-diamidino-2-phenylindole, dihydrochloride (DAPI, DNA). (**B**) Quantification of average spindle-localized F-actin in **A**. (**C**) Representative confocal images of a metaphase I oocyte expressing mRuby fused to full-length mouse myosin VIIb (mRuby-MyoVIIb, pseudo-cyan) immunostained with pericentrin antibody (pMTOCs, magenta). (**D**) Representative confocal images of metaphase I oocytes microinjected with MyoX siRNA or MyoX siRNA + MyoVIIb cRNA at the germinal vesicle stage. The oocytes were labeled with phalloidin (F-actin, gray) and pericentrin (pMTOCs, magenta). DNA was labeled with DAPI (blue). Scale bar represents 10 μm. (**E**) Plot profile of pMTOCs (pericentrin fluorescence) along the longitudinal axis of the spindle in the groups indicted in **D**. (**F**) Quantification of pMTOC volume at the spindle poles to the total volume of all pMTOCs in the spindle. (**G**) Average pMTOC number quantification after 3D reconstruction of pericentrin from oocytes in **D**. (**H**) Average pMTOC volume quantification after 3D reconstruction of pericentrin from oocytes in **D**. (**I**) Average spindle-localized F-actin fluorescence intensity of oocytes indicated in **D**. (**J**) Average cytoplasmic F-actin fluorescence intensity of oocytes indicated in **D**. Data are represented as means ± SEM. Asterisks represent significant differences, ** p<0.01, *** p<0.001, and **** p<0.0001. The total number of analyzed oocytes per each experimental group (from 3 independent replicates) is specified above each graph.

### Spindle-localized F-actin dynamics are critical for pMTOC sorting and clustering

To further confirm that spindle-localized F-actin (and its dynamics) is required for pMTOC sorting and clustering, we optimized the protocol of a novel cell-permeable jasplakinolide-derived compound, Optojasp-1. The F-actin stalling ligand 2 was modified by adding an azobenzene photoswitch that can be activated spatiotemporally by 405 nm light^70^ (Fig. 5A). Thus, Optojasp-1 binds to F-actin in the entire cell, while it is still inactive (i.e., not perturbing F-actin dynamics). Only upon 405 nm laser exposure, the photoswitchable Optojap-1 undergoes structural conformation and becomes active (i.e., perturbing F-actin dynamics). This molecule, Optojasp-1, can be used to inhibit F-actin dynamics in a desired region of interest for approximately 4 h before the compound returns to its inactive state^70, 71^, thus offering high spatial and temporal resolution. Indeed, in contrast to control oocytes, Optojasp-treated oocytes exposed to a 405 nm laser at a single spindle pole perturbed F-actin dynamics, at the exposed spindle pole but not at the opposite spindle pole (Fig. 5B,C and Supplementary Movie 2) or near chromosomes (Supplementary Fig. 10A-C). It is of note that exposing one spindle pole to 405 nm laser (without Optojasp-1 treatment) had no effect on spindle pole-localized F-actin localization or dynamics (Supplementary Fig. 10D,E). Moreover, selective disruption of F-actin dynamics only at the spindle poles using Optojasp approach had no effect on K-fiber stability (which is likely regulated by F-actin localization near chromosomes) (Supplementary Fig. 10F,G). Thus, our results indicate that Optojasp-1 is an efficient tool to locally disrupt F-actin dynamics in our system. We did not observe excessive disordered F-actin nucleation (Fig. 5B) or spindle-localized G-actin reduction in Optojasp-treated spindle-exposed oocytes (Supplementary Fig. 10H,I) when compared to controls. Thus, Optojasp-1, at the used concentration, does not exhaust the G-actin pool, but stabilizes F-actin (perturbs its dynamics) and promotes thick F-actin bundles and aggregates, leading ultimately to disrupted F-actin network (i.e., disrupted spindle-localized F-actin structure, Supplementary Fig. 10J). Using high-resolution time-lapse confocal microscopy, we tracked the numbers and volumes of fluorescently labeled pMTOCs (Cep192-mCherry) and the spindle (SiR-tubulin) in Optojasp-treated oocytes in which the spindle was exposed to 405 nm laser 1 h post-NEBD to perturb spindle-localized F-actin dynamics. To exclude the possibility of any optojasp/light-related off-target effect on our result interpretation, we used two control groups: non-Optojasp-treated oocytes were exposed to the 405 nm laser at the spindle and Optojasp-treated oocytes were exposed to the 405 nm laser in the cytoplasm to perturb F-actin outside the spindle (Fig. 5D,E). In contrast to control oocytes, disrupting F-actin dynamics at the spindle, but not in the cytoplasm, prevented proper pMTOC sorting and clustering (Fig. 5E-G), further confirming that spindle-localized F-actin is required to properly regulate pMTOC sorting and clustering during oocyte meiosis.

**Figure 5:**
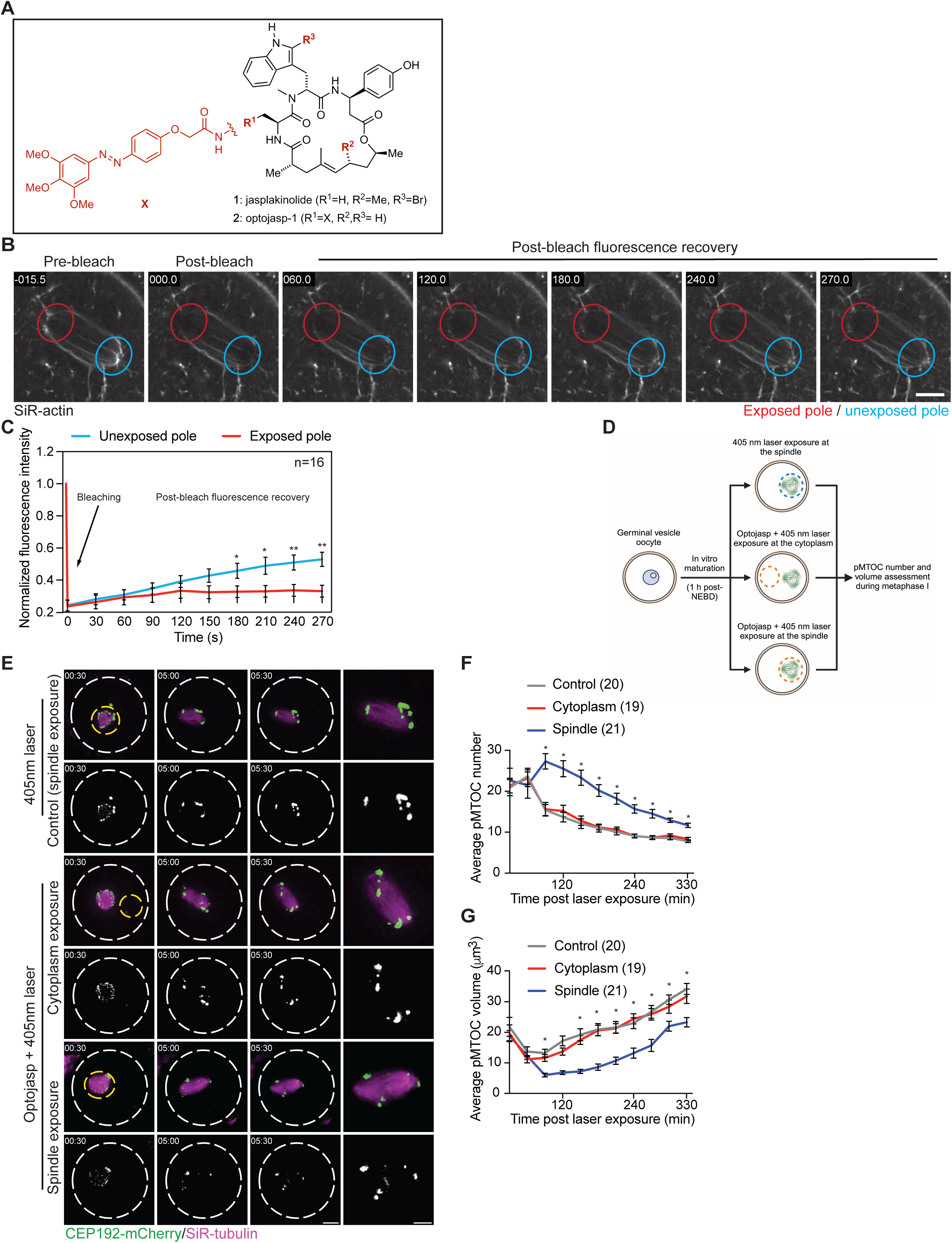
Spindle-localized F-actin regulates pMTOC sorting and clustering. (**A**) Chemical structure of Optojasp-1. (**B**) Metaphase I oocytes expressing CEP192-mCherry and labeled with SiR-actin (F-actin) were treated with Optojasp-1 and exposed to 405 nm laser at one spindle pole (spindle pole-localized F-actin dynamic perturbation). Bleaching was performed 1 h post-405 nm laser exposure at the two spindle poles using 100% 647 nm laser power (100 ms laser pulse) followed by imaging for an additional 5 minutes to monitor the fluorescence recovery after photobleaching (FRAP). (**C**) Quantification of SiR-actin fluorescence recovery after photobleaching (FRAP). (**D**) Schematic diagram shows the experimental design of selectively disrupting spindle-localized F-actin using Optojasp-1. (**E**) Time-lapse live imaging confocal microscopy of pMTOCs (Cep192-mCherry; pseudo green) and SiR-tubulin (spindle; magenta) in prometaphase I mouse oocytes treated with DMSO and exposed to 405 nm laser at the spindle (control 1), treated with Optojasp-1 and exposed to 405 nm laser at the cytoplasm (control 2), or treated with Optojasp-1 and exposed to 405 nm laser at the spindle (spindle-localized F-actin perturbation). Images were taken every 30 min. Scale bar represents 20 μm. (**F**) Average pMTOC number quantification after 3D reconstruction of pMTOCs for the oocytes illustrated in **E**. (**G**) Average pMTOC volume quantification after 3D reconstruction of pMTOCs for the oocytes illustrated in **E**. Data are represented as means ± SEM. Asterisk represents significant differences, * p<0.05 and ** p<0.01. The total number of analyzed oocytes per each experimental group (from 3 independent replicates) is specified above each graph.

### MyoX and F-actin organize pMTOCs through a positive feedback mechanism

Our results revealed that spindle-localized F-actin regulates pMTOC sorting and clustering. However, the underlying molecular mechanism of F-actin-mediated pMTOC sorting and clustering is unknown. We hypothesized that MyoX recruits F-actin to the MT spindle which, in turn, forms the platform for the F-actin-based motor MyoX to move MTOCs towards the spindle pole, leading to their clustering and sorting. MyoX is an F-actin-based motor that moves toward F-actin barbed ends. To examine the orientation of spindle-localized F-actin, we employed a FRAP assay to assess the direction of F-actin growth during recovery. Approximately 90% of F-actin fibers grew towards the spindle poles, indicating that the barbed ends of spindle-localized F-actin are oriented toward the spindle poles (Supplementary Fig. 11A,B and Supplementary Movie 3). This is expected because regardless of F-actin’s barbed end direction it grows until reaching the spindle pole, whether the nearest spindle pole or the opposite spindle pole. Indeed, an F-actin fiber near one spindle pole can grow towards and reach the opposite spindle pole, making some F-actin fibers longer and appear as they span the entire spindle length (Supplementary Fig. 10D). To investigate whether MyoX moves MTOCs along F-actin fibers, we expressed mRuby-tagged versions (Fig. 6A) of full-length mouse MyoX; MyTH4-mutant MyoX, which lacks MT binding ability; or headless MH-mutant MyoX, which localizes to MTs but lacks the motor head activity^72, 73^. Expectedly, the headless MH-mutant MyoX exhibited reduced motility on actin fibers, compared to full-length mouse MyoX (Supplementary Fig. 11C,D). The minimal movement observed in headless MH-mutant MyoX could be due to its tethering to an endogenous MyoX monomer. We then assessed 1) MyoX’s localization to the spindle, 2) localization of F-actin to the spindle, and 3) pMTOC sorting and clustering. We found that oocytes expressing full-length MyoX had normal MyoX localization (mRuby at the spindle and pMTOCs, Fig. 6B and Supplementary Fig. 11E), typical spindle-localized F-actin (Fig. 6B,C) and proper pMTOC sorting (Fig. 6B,D,E) and clustering (Fig. 6B,F,G). Expression of the MyTH4 dominant-negative MyoX mutant decreased mRuby MyoX at the spindle (Fig. 6B and Supplementary Fig. 11E) and reduced F-actin localization to the spindle (Fig. 6B,C) without perturbing cytoplasmic F-actin (Supplementary Fig. 11F), confirming that F-actin recruitment to the spindle depends on MyoX binding to MTs, and resulted in pMTOC clustering and sorting defects (Fig. 6B,D-G). Importantly, compared to controls, expressing the headless MyoX mutant, which lacks motor activity, decreased the average speed of pMTOC movement towards the spindle poles (Supplementary Fig. 11G,H and Supplementary Movie 4 and Movie 5), resulting in pMTOC clustering and sorting defects (Fig. 6B,D-G) even in the presence of normal MyoX localization (through the MyTH4 domain) and F-actin localization to the spindle (most likely occurs through its FERM domain^49, 50, 58^ that has been shown to bind F-actin in vitro) (Fig. 6B,C and Supplementary Fig. 11E). Together, these results indicate that the motor activity of MyoX contributes to pMTOC sorting and clustering and suggest some positive feedback mechanism between F-actin and MyoX. It suggests a model in which (1) MyoX localizes to the spindle and pMTOCs, (2) MyoX recruits F-actin to the spindle and pMTOCs, and (3) F-actin provides the platform for F-actin based motors, at least MyoX, which attach to unclustered and unsorted pMTOCs to move them along F-actin fibers towards the spindle poles (i.e., towards the barbed ends of F-actin), leading to their clustering and sorting (Fig. 6H).

**Figure 6:**
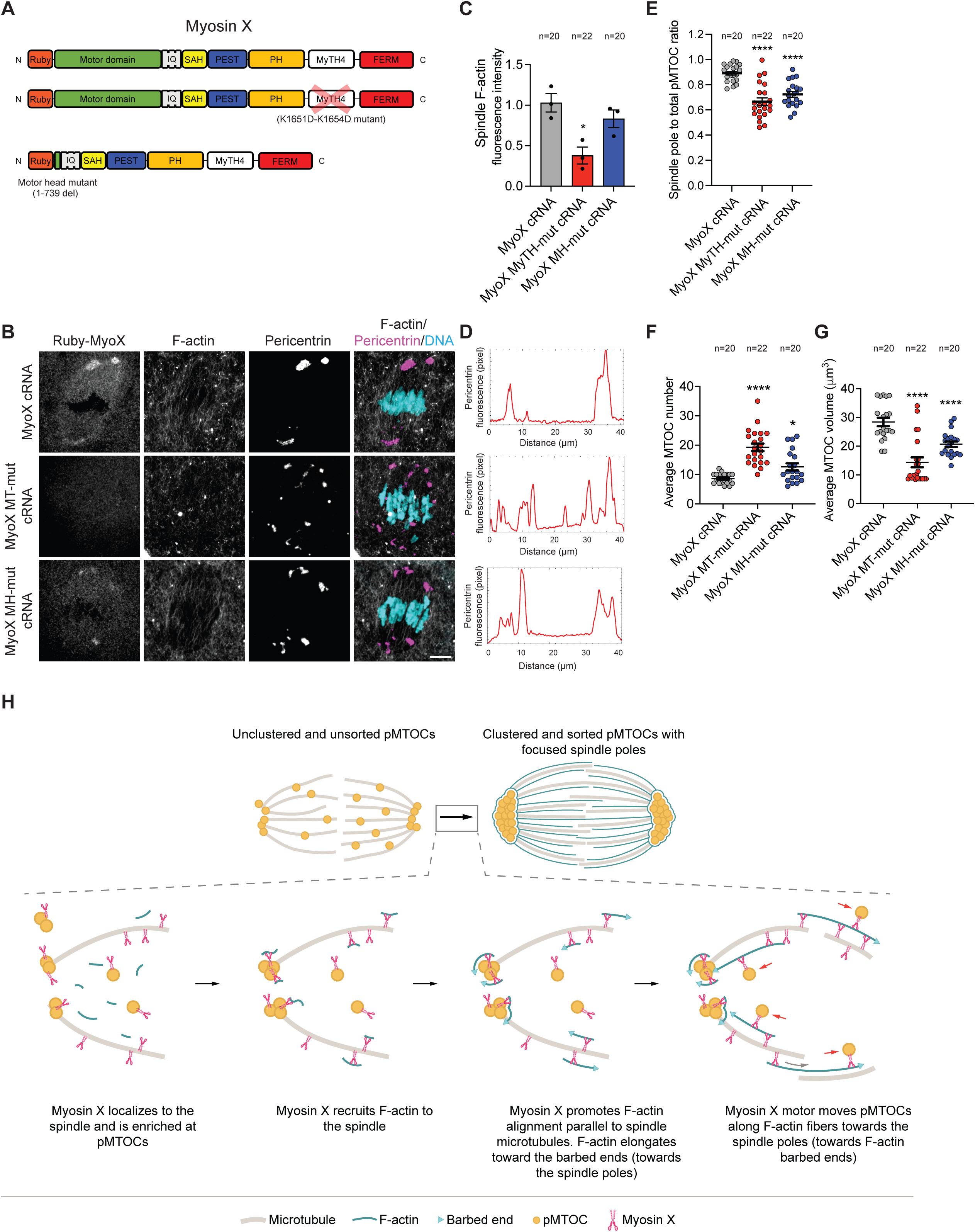
Positive feedback mechanism between Myosin X and F-actin is required for efficient pMTOC sorting and clustering. (**A**) Schematic diagrams showing the domains of Myosin X constructs: full length Myosin X (upper), microtubule binding domain (MyTH4, K1651D/K1654D) Myosin X mutant (MyoX MyTH-mut, middle), and headless Myosin X mutant (lacking the motor head, 1-739 deletion, MyoX MH-mut, lower). (**B**) Representative confocal images of metaphase I oocytes microinjected with mRuby-tagged versions of MyoX, MyoX MyTH-mut, or MyoX MH-mut cRNA at the germinal vesicle stage. The oocytes were labeled with phalloidin (F-actin, gray) and pricentrin antibody (magenta). DNA was labeled with 4′,6-diamidino-2-phenylindole, dihydrochloride (DAPI, cyan). Scale bar represents 10 μm. (**C**) Average spindle-localized F-actin fluorescence intensity of oocytes microinjected with MyoX, MyoX MyTH-mut, or MyoX MH-mut cRNA. (**D**) Plot profile of pMTOCs (pericentrin fluorescence) along the longitudinal axis of the spindle in the groups indicted in **B**. (**E**) Quantification of pMTOC volume at the spindle poles to the total volume of all pMTOCs in the spindle. (**F**) Average pMTOC number quantification after 3D reconstruction of pericentrin from oocytes in **B**. (**G**) Average pMTOC volume quantification after 3D reconstruction of pericentrin from oocytes in **B**. Data are represented as means ± SEM. Asterisks represent significant differences, * p<0.05, ** p<0.01, and **** p<0.0001. The total number of analyzed oocytes per each experimental group (from 3 independent replicates) is specified above each graph. (**H**) Schematic diagram depicting the positive feedback mechanism by which pMTOCs are clustered, sorted, and tethered/maintained.

### Spindle pole-localized F-actin maintains clustered pMTOCs to prevent chromosome missegregation

Time-lapse tracking of pMTOCs in Cep192-eGFP reporter oocytes revealed that, after pMTOC clustering and sorting at early metaphase I, pMTOCs were maintained at the spindle poles for an additional 1-2 h before chromosome segregation (Fig 2A). Therefore, we hypothesized that a third step of pMTOC organization is required to ensure faithful chromosome segregation, in which sorted and clustered pMTOCs are tethered/maintained at spindle poles until anaphase I onset. Previous studies showed that F-actin forms a cage surrounding the meiotic spindle^24, 25, 28^. Using super-resolution microscopy, we observed a previously undocumented F-actin cage-like structure surrounding the already sorted and clustered pMTOCs at metaphase I in 78.33% of oocytes (Fig. 7A,B). This observation led us to hypothesize that F-actin not only plays a role in pMTOC sorting and clustering but is also required to maintain/tether already clustered and sorted pMTOCs at the spindle poles. To investigate whether F-actin is required to maintain pMTOC sorting and clustering, we allowed oocytes to mature in vitro to metaphase I (5 h post-NEBD, the stage at which pMTOCs are already sorted and clustered) and then treated oocytes with DMSO or CytoD for an additional 2 hours (7 h-post NEBD) before assessing pMTOC sorting and clustering. Indeed, at 5 h post-NEBD, pMTOCs were largely sorted and clustered (Fig. 7C-G). Strikingly, in contrast to DMSO-treated oocytes (7 h-post NEBD), oocytes with perturbed F-actin showed increased numbers of pMTOCs with decreased volumes (pMTOC clustering defect; Fig. 7C,F,G), that were inappropriately scattered along the spindle axis with decreased spindle pole to total pMTOC volume (pMTOC sorting defect; Fig. 7C-E). These findings indicate that F-actin is required to maintain and tether the already clustered and sorted pMTOCs. To probe the significance of the spindle pole-localized F-actin cage-like structure in the maintenance of pMTOC sorting and clustering, we selectively perturbed the dynamics of spindle pole-localized F-actin using Optojasp-1 and assessed pMTOC behavior using time-lapse confocal microscopy. Small circular areas surrounding only one or both spindle poles were marked and exposed to 405 nm light in Optojasp-treated oocytes at metaphase I (5 h post-NEBD, when pMTOCs are already sorted and clustered) (Fig. 7H). Control Optojasp-treated oocytes were exposed to the same protocol except random areas of the cytoplasm, immediately adjacent to and equal in size to the spindle pole regions, were exposed to 405 nm light. In accord with the effect of CytoD on pMTOC sorting and clustering maintenance, the selective disruption of F-actin dynamics at both spindle poles resulted in pMTOC fragmentation and scattering along the spindle axis (Fig. 7H-J) when compared to control (cytoplasmic exposure). Because spindle-localized F-actin arises from cytoplasmic F-actin, it is plausible to anticipate a more pronounced effect on pMTOC fragmentation in CytoD-treated oocytes, in which cytoplasmic F-actin meshwork is also disrupted, compared to Optojasp-1-mediated F-actin dynamic perturbation at the spindle pole. Importantly, when F-actin was disrupted at a single spindle pole, the exposed pole showed pMTOC fragmentation and scattering, while the unexposed pole remained intact (Fig. 7H-J). This suggests that the F-actin cage-like enrichment at the spindle poles is a dynamic structure that is required to maintain the sorting and clustering of pMTOCs during oocyte meiosis.

**Figure 7:**
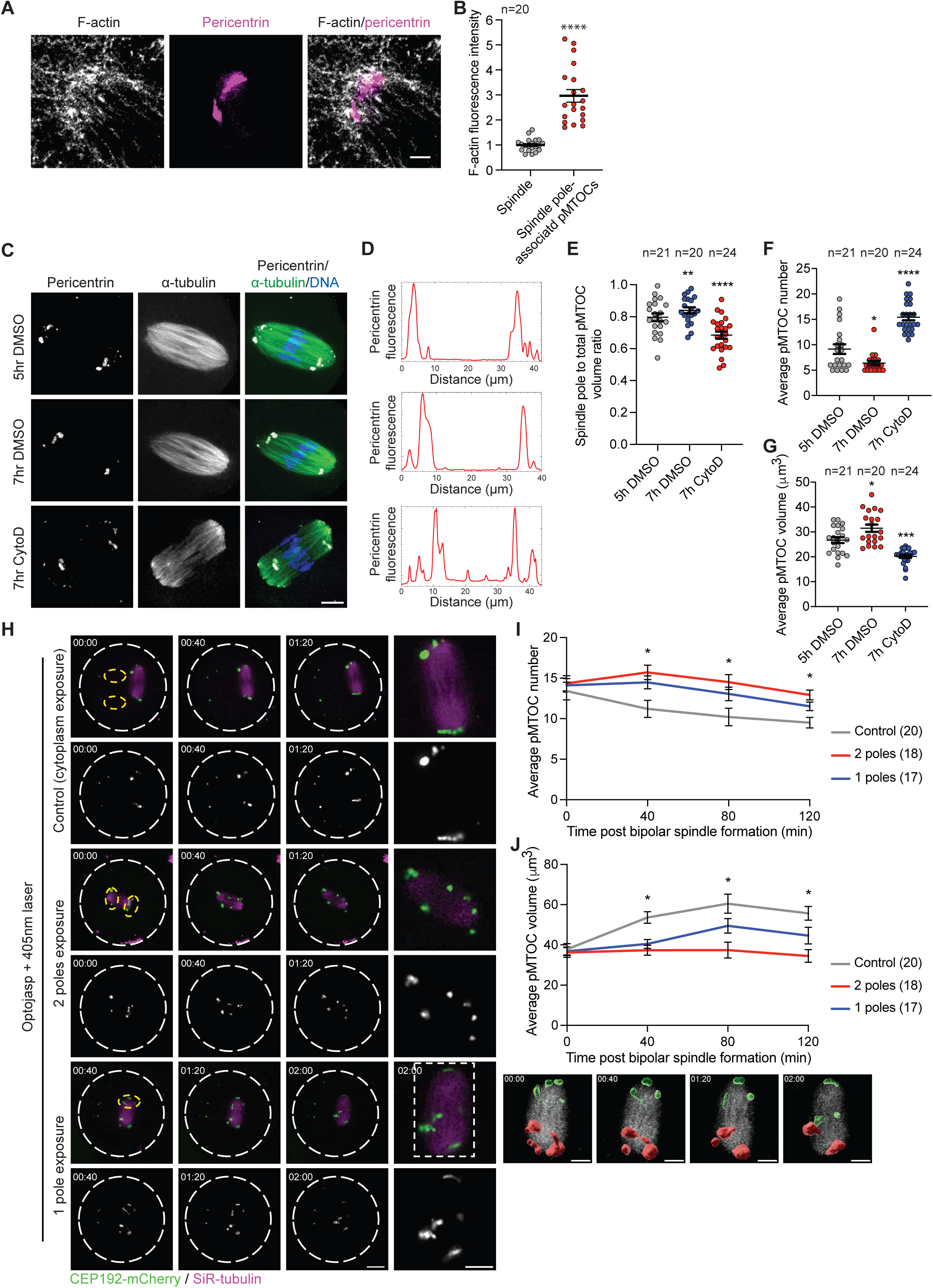
Spindle pole-localized F-actin is required to regulate pMTOC sorting and clustering maintenance. (**A**) Representative Super-resolution Stimulated Emission Depletion (STED) confocal image of F-actin cage-like enrichment at the spindle poles in a metaphase I oocyte. The oocyte was labeled with phalloidin (F-actin, gray) and pricentrin antibody (magenta). Scale bar represents 10 μm. (**B**) Quantification of F-actin enrichment at pMTOCs. (**C**) Representative confocal images of metaphase I oocytes treated with DMSO or CytoD at early metaphase I (5h post-NEBD) and fixed at late metaphase (7h post-NEBD). The oocytes were immunostained with α-tubulin to label the spindle (green) and pericentrin to label MTOCs (gray). DNA was labeled with 4′,6- diamidino-2-phenylindole, dihydrochloride (DAPI, blue). Scale bar represents 10 μm. (**D**) Plot profile of pMTOCs (pericentrin fluorescence) along the longitudinal axis of the spindle in the groups indicted in **C**. (**E**) Quantification of pMTOC volume at the spindle poles to the total volume of all pMTOCs in the spindle. (**F**) Average pMTOC number quantification after 3D reconstruction of pericentrin from oocytes in **C**. (**G**) Average pMTOC volume quantification after 3D reconstruction of pericentrin from oocytes in **C**. (**H**) Time-lapse live imaging confocal microscopy of pMTOCs (Cep192-mCherry; pseudo green) and SiR-tubulin (spindle; magenta) in metaphase I mouse oocytes treated with Optojasp-1 and exposed to 405 nm laser at the cytoplasm (control) or treated with Optojasp-1 and exposed to 405 nm laser at one, or at both spindle poles. Images were taken every 40 min. Scale bar represents 20 μm. Right panels show 3D reconstruction of pMTOCs for the oocytes treated with Optojasp-1 and exposed to 405 nm laser at one spindle pole. Exposed pMTOCs are labeled in green, whereas non-exposed pMTOCs are labeled in Red. (**I**) Average pMTOC number quantification after 3D reconstruction of pMTOCs for the oocytes illustrated in **H**. (**J**) Average pMTOC volume quantification after 3D reconstruction of pMTOCs for the oocytes illustrated in **H**. Data are represented as means ± SEM. Asterisk represents significant differences, * p<0.05, *** p<0.001 and **** p<0.0001. The total number of analyzed oocytes per each experimental group (from 3 independent replicates) is specified above each graph.

Chromosome segregation errors occur frequently during oocyte meiosis, especially meiosis I. In all previous F-actin perturbations, including the expression of MyoX MyTH4-mutant or MyoX headless MH-mutant (Supplementary Fig. 12A-C), we observed a significant increase in the incidence of chromosome misalignment and lagging chromosomes when compared to control oocytes. To determine whether spindle pole-localized F-actin is essential to ensure proper chromosome segregation, we selectively perturbed spindle pole-localized F-actin at the two spindle poles using Optojasp-mediated F-actin perturbation followed by time-lapse confocal microscopy to assess chromosome segregation. Disrupting F-actin dynamics at both spindle poles increased the percentage of chromosome misalignment at metaphase I (Supplementary Fig. 12D,E), the incidence of lagging chromosomes (> five-fold) during anaphase I/telophase I (Supplementary Fig. 12D,F) and increased the incidence of aneuploidy at metaphase II (Supplementary Fig. 12G,H) when compared to control oocytes. Taken together, these findings indicate that spindle-localized F-actin, is required for efficient pMTOC sorting, clustering and maintenance at the spindle poles during oocyte meiosis and to prevent the development of aneuploid gametes.

## Discussion

Recently, F-actin has been linked to various processes in oocyte meiosis^20, 21, 23–26, 28^. While the localization of F-actin at the spindle in mammalian oocytes is established ^24, 25, 28, 29^, the precise mechanism underlying this localization remains unknown partly because of a dearth of specialized tools for selectively disrupting spindle-localized F-actin in mammalian oocytes. We have found that F-actin localization to the spindle MTs relies on unconventional myosins, MyoX and MyoVIIb. By knocking down MyoX/MyoVIIb or by utilizing an optimized photoswitchable jasplakinolide, Optojasp-1, we selectively disrupted spindle-localized F-actin or polymerization dynamics to reveal a novel role for F-actin in regulating pMTOC clustering and sorting, processes critical for ensuring the fidelity of meiotic spindle assembly. We also observed an F-actin cage-like enrichment surrounding pMTOCs that were already clustered and sorted. We found that the dynamic behavior of this spindle pole-localized F-actin cage-like enrichment is essential for maintaining pMTOC clustering and sorting. A previous study demonstrated that F-actin is required for proper chromosome segregation by promoting K-fiber stability^28^. In our study, we demonstrated that perturbing the spindle pole-localized F-actin during metaphase I resulted in pMTOC clustering and sorting defects and increased the incidence of chromosome segregation errors, independent of F-actin role on K-fiber stability. Our results suggest that F-actin at the spindle plays two distinct roles to ensure faithful chromosome segregation. F-actin localization at the spindle poles (as shown in this study) regulates MTOC clustering and sorting, whereas F-actin localization at the chromosomes promotes K-fiber stability before anaphase I^28^.

The long-standing view has been that MTs (and their associated proteins) are the sole cytoskeletal components responsible for proper spindle assembly in all cell types^13, 18, 19^. The unexpected role of spindle-localized F-actin in regulating pMTOC organization might provide an explanation to the abnormal spindle morphology associated with the perturbation of F-actin/actin nucleators in mammalian oocytes^29, 74–78^. We speculate that, unlike somatic cell spindles, the spindle in mammalian oocytes is large and relies on numerous acentriolar MTOCs, a situation that may be too complex for MTs alone to efficiently cluster and sort multiple MTOCs into two poles. In addition to the established role of MTs and their associated proteins in pMTOC clustering and spindle pole focusing, we uncovered a novel role of spindle-localized F-actin in regulating these processes. Understanding how the two major cytoskeletal components, MTs and F-actin, and their regulatory molecules interact at the molecular level to assemble meiotic spindles is a key focus of our future studies.

While there is recent evidence demonstrating F-actin localization at centrosomes and the mitotic spindle in somatic cells^30, 31, 79, 80^, its specific contribution to the bipolar spindle formation remains uncertain. We found that F-actin localizes to the spindle and pMTOCs in mouse oocytes, and this localization is critical for pMTOC clustering and sorting. Our data suggest a model in which MyoX first localizes to the spindle and pMTOCs through its MT binding domain. Given that MyoX has both F-actin and MT binding domains, it then recruits F-actin to spindle MTs and promotes its alignment along spindle MT fibers, essential steps for the proper assembly of spindle-localized F-actin and pMTOC-associated F-actin. Because MyoX attaches to all pMTOCs, it moves non-clustered pMTOCs along the already assembled actin filaments towards the spindle poles (i.e., towards F-actin barbed ends), driven by its motor activity, promoting pMTOC clustering and sorting. Moreover, our high-resolution time-lapse imaging of F-actin and pMTOCs revealed that, throughout meiotic progression, F-actin is increasingly recruited to pMTOCs. Therefore, it is plausible that F-actin recruitment to large pMTOCs while it is still attached to smaller pMTOCs also contributes to pMTOC clustering and sorting.

A specific family of unconventional myosins, F-actin-based motors, contains MyTH4 domains, which mediate MT binding^49, 50^. Such MyTH4-containing unconventional MyTH4 myosins, that include MyoX and MyoVIIb, may serve as bridging elements, recruiting F-actin to the spindle MTs. We found that disrupting MyoX or MyoVIIb, but not MyoXVa, decreased F-actin localization at the spindle, leading to pMTOC clustering and sorting defects. These findings suggest that MyoX and MyoVIIb share similar roles in regulating F-actin localization at the spindle and their levels “combined” are required to carry out this function. Notably, ectopic expression of MyoVIIb significantly rescued both the localization of F-actin at the spindle and the pMTOC clustering and sorting defects in MyoX KD oocytes. The hypothesis that MyoX, MyoVIIb and potentially other unidentified regulators have overlapping functions is supported by the finding that MyoX knockout mice display normal fertility, oocyte spindle morphology and mitosis^63, 64^, despite several *in vitro* studies showing MyoX to be indispensable for meiotic and mitotic divisions^51, 52, 55^. This suggests the presence of compensatory mechanisms for the localization of F-actin at the spindle in MyoX knockout oocytes. Such compensatory functions among related proteins are well-documented in genetic knockout models but are less likely to occur in knockdown systems. Unlike knockout mice, which allow a relatively long window (several weeks) for transcriptional or post-transcriptional compensation, knockdown approaches offer only a short timeframe (a few hours) for such adaptation. This limitation is further exacerbated in fully grown oocytes, which are transcriptionally silent, limiting their ability to timely upregulate compensatory proteins in response to knockdown. It is of future interest to investigate whether F-actin localization at the spindle is compensated in MyoX knockout oocytes and whether unconventional myosins or MT-associated proteins, which have the ability to cross-link MTs and F-actin in mitotic cells^49, 50, 58^, contribute to this phenomenon.

The presence of extra centrosomes in mitotic cells can result in multipolar cell division, leading to impaired cell viability^81^. Similar to mouse oocytes, cancer cells have evolved unique mechanisms to cluster these extra centrosomes into two spindle poles, thereby preventing multipolar cell divisions and ensuring their survival^54^. A growing body of evidence suggests that F-actin and its associated proteins play a role in regulating centrosome clustering in cells that possess supernumerary centrosomes. Disrupting F-actin in both mammalian cancer cell lines (N1E-115)^54^ or mouse oocytes (this study) can result in centrosome/pMTOC clustering defects. The involvement of F-actin in regulating centrosome clustering in cancer cells appears to be independent of intrinsic spindle force^54^. In contrast, mouse oocytes appear to rely on spindle-localized F-actin, rather than cortical F-actin, for regulating pMTOC clustering. Despite potential differences in the underlying mechanisms, the involvement of F-actin in centrosome/pMTOC clustering seems to be conserved in cells harboring supernumerary centrosomes/MTOCs. In human oocytes, perinuclear MTOCs (huoMTOCs) undergo fragmentation followed by their sorting to kinetochores for the initiation of bipolar spindle assembly^10^. F-actin localizes to the human oocyte spindle and its disruption leads to spindle defects. Therefore, it is tempting to speculate that F-actin also regulates huoMTOC sorting in human oocytes, ensuring the correct assembly of bipolar spindles— a critical process for accurate chromosome segregation.

## Methods

### Experimental mice

All experiments and mice care were conducting according to the regulations of the University of Missouri (Animal Care Quality Assurance Ref. Number, 9695). CF-1 mice were purchased from Envigo, Indianapolis, IN, USA. Unless otherwise specified, all experiments were conducted using CF-1 mice. Cep192-eGfp reporter mice were previously generated^16^. Homozygous Cep192-eGfp breeding pairs were used to maintain the colony. All animals had ad libitum access to water and food at 21 °C with 55% humidity and in 12-hour light/dark cycle.

### Cloning and in vitro cRNA synthesis

pRN3P-mRuby-MyoVIIb, pRN3-mRuby-MyoX, pRN3-mRuby-MyoX (1-739 deletion) constructs were generated by cloning mMyoVIIb, mMyoX and mMyoX 1-739 deletion (GenScript Biotech, Piscataway, New Jersey, USA) in frame with mRuby into pRN3P vector. mMyoX K1651D/K1654D mutant was generated by site-directed mutagenesis using the QuikChange Multi-site Mutagenesis kit (Agilent Technologies) according to the manufacturer’s instructions. MyoX and MyoVIIb siRNA resistant mutants were generated by GenScript Biotech. Generation of *eGfp-Eb3* was described previously^15^. DNA linearization was carried out using SFiI restriction enzyme (New England BioLabs) followed by DNA purification according to the manufacturer’s protocol (QIAquick PCR Purification, Qiagen). In vitro transcription of purified DNA was done using an mMessage mMachine T3 kit (Ambion), following the manufacturer’s instructions. Purification of cRNA was carried out using an RNAEasy kit (Qiagen) and stored at −80°C.

### Oocyte collection, microinjection and culture

Full-grown GV-stage-arrested mouse oocytes were collected from 6-8-week-old CF-1 or Cep192-eGfp reporter CF-1 female mice. Cumulus oocyte complexes (COCs) were collected in bicarbonate-free minimal essential medium (MEM) supplemented with 3 mg/mL polyvinylpyrolidone (PVP) and 25 mM Hepes (pH 7.3). The oocytes were denuded from cumulus cells by mechanical pipetting prior to oocyte maturation in Chatot, Ziomek, and Bavister (CZB) medium^82^ at 37°C in a humidified atmosphere containing 5% CO_2_. Denuded GV oocytes were microinjected with 5 pl of the indicated cRNAs or siRNAs using Eppendorf FemtoJet 4i. The microinjection medium was MEM-PVP-Hepes medium supplemented with 2.5 μM milrinone (MilliporeSigma, St. Louis, MO, USA, M4659), a phospho diesterase inhibitor to prevent the meiotic resumption^83^. The injected oocytes were cultured for 16 h in CZB medium supplemented with 2.5 μM milrinone at 37°C in a humidified atmosphere containing 5% CO_2_. The oocytes were then allowed to resume meiosis in milrinone-free CZB medium. Unless otherwise specified, metaphase I stage oocytes were collected 5 h after NEBD (∼7 h after maturation). Oocytes that did not progress beyond NEBD were excluded from experiments.

Porcine and bovine ovaries were collected at the slaughterhouse and transported to the laboratory at the University of Missouri within 2 h and 4 h, respectively. The cumulus oocyte complexes (COCs) were aspirated from 3-8 mm follicles using 10 mL syringe equipped with an 18-gauge needle. Only good quality oocytes, with homogenous cytoplasm and surrounded by more than 3 layers of cumulus cells, were selected. The COCs were cultured in chemically defined maturation medium, TCM 199 (Invitrogen, Grand Island, NY), supplemented with 10% bovine calf serum, 1 mM L-glutamine, 0.2 mM sodium pyruvate, 20 μg/mL follicle stimulating hormone [FSH; Folltropin-V, Vetoquinol, TX, USA], 2 μg/mL β-estradiol and 10 μg/mL gentamycin at 38. 5°C and 5% CO_2_ in humidified air. Deidentified immature human COCs were received from the University of Missouri OneHealth Biorepository and Center for Assisted Reproduction, Department of Gynaecology, Obstetrics and Neonatology, First Faculty of Medicine, Charles University and General University Hospital in Prague, Czech Republic. The immature human COCs were retrieved during stimulated ovulation cycles with the patients’ consent and ethical committee approval (no. 1459/23 IS, D; BIOCEV 012019). Cumulus cells were removed mechanically by gentle pipetting and the oocytes were matured individually in 20 μl drops of G-MOPS medium (Vitrolife, 10129) at 37.2 °C and ambient atmosphere or Sequential Fert medium (Origio, 83020010A) at 37.2 °C and 6.2% CO_2_ and 5% O2. The maturation medium drops were overlayed with 500 μl of Ovoil (Vitrolife, 10029). The maturation status and the meiotic spindle were regularly monitored using Spindle visualization with Olympus IX3 – ICSI/IMSI system. CytoD was added to the maturation medium at 5 h post-NEBD. The oocytes were allowed to mature in CytoD-containing medium for an additional 5 h.

### Drug treatments

Cytochalasin D (5 μg/mL, MilliporeSigma, C2618), Latrunculin B (5 μM, MilliporeSigma, L5288), Jasplakinolide (0.01 μM, Invitrogen, J7473), CK-666 (50 μM, MilliporeSigma, 182515), SMIFH2 (300 μM, MilliporeSigma, S4826), nocodazole (5 μM, MilliporeSigma, M1404), MG-132 (7.5 μM, MilliporeSigma, 474790) or blebbistatin (100 μM, MilliporeSigma, B0560) were dissolved in DMSO (MilliporeSigma, D2650) and added to the maturation medium at NEBD. During time-lapse live imaging, a final concentration of 100 nM of SiR-tubulin (Cytoskeleton, CY-SC002), SiR-actin (Cytoskeleton, CY-SC001) or SiR-DNA (Cytoskeleton, CY-SC007) was added to the culture medium to label MTs, F-actin and DNA, respectively.

### Optojasp-mediated F-actin disruption

Optojasp-1, the first-generation cell permeable photoswitchable compound synthetized from Jasplakinolide^70^. Optojasp-1 itself remains inactive until it is activated upon exposure to 405 nm light, resulting in F-actin dynamics perturbation^70^. To perturb F-actin/dynamics, Optojasp-1 was added to the culture medium at NEBD at a concentration of 0.01 μM^70^. To activate Optojasp-1 and disrupt F-actin selectively at the spindle/spindle poles, 405 nm wavelength laser was exposed to a specific region of interest surrounding the spindle/spindle poles followed by time-lapse confocal microscopy to track the fluorescently labeled spindle (SiR-tubulin) and pMTOCs (mCherry-Cep192). Non-Optojasp-treated oocytes in which the spindle or the spindle poles were exposed to 405 nm laser and Optojasp-treated oocytes in which an area of the cytoplasm (adjacent to and of equal size to the spindle/spindle poles) was exposed to 405 nm laser exposure served as control groups.

### Immunocytochemistry and confocal microscopy

Mouse and bovine oocytes were fixed for 1 h at room temperature in freshly prepared 3.7% paraformaldehyde solution (MilliporeSigma, P6148) dissolved in phosphate buffer saline (PBS). Fixed mouse and porcine oocytes were permeabilized for 20 min at room temperature in PBS containing 0.1% Triton X-100. Permeabilized oocytes were incubated for 30 min at room temperature in PBS containing 0.3% BSA and 0.01% Tween-20 (blocking solution). Human and porcine oocytes were fixed and permeabilized in 100 mM PIPES (AppliChem, A1079) solution (pH 7.0) containing 5 mM magnesium chloride hexahydrate, 2 mM EGTA, 2% formaldehyde (MilliporeSigma, F1635), 0.5% Triton X-100, and 10 nM paclitaxel (MilliporeSigma, T7191) for 30 min at 37 °C, as previously described^29^. For primary antibody incubation, oocytes were incubated for 1 hour at room temperature in the indicated primary antibodies, followed by three 9-min washes in blocking solution. Similarly, secondary antibodies incubation was carried out by incubating oocytes for 1 hour at room temperature, followed by three 9-min washes in blocking solution. Texas Red-X Phalloidin (1:50, Invitrogen, T7471), conjugated Alexa Fluor 488-Phalloidin (1:50, Invitrogen, A12379) or Abberior Star 635P (1:50, Abberior GmhB, ST635P-0100-20UG) for STED was added to the secondary antibody solution to detect F-actin. G-actin was stained using Fluorescent Deoxyribonuclease I (DNase I) conjugate (10 μg/ml, Molecular Probes, D12372) according to^84^. Immunolabeled oocytes were mounted on glass slides with VECTASHIELD containing 4’,6-Diamidino-2-Phenylindole, Dihydrochloride (DAPI; Vector Laboratories, Burlingame, CA, USA) to label DNA. Hoechst 33342 (5 mg/ml, Molecular Probes H3570) was used to label the DNA and a hardening mounting medium AD-MOUNT H (ADVI, Ricany, CZ) was applied for STED super-resolution imaging. Fluorescence images were taken using a 100x oil objective of Leica Stellaris 5 confocal microscope. Super-resolution images were detected under a 63X objective using Leica TCS SP8 confocal microscope equipped with 3-color, 3-D STED super-resolution 3X system or Leica TCS SP8 STED 3X microscope equipped with a pulsed white light laser (WLL2) for excitation and a pulsed 775 nm laser for emission depletion (Leica Microsystems).

Oocytes were imaged using 0.5 or 1 μm Z-intervals spanning the meiotic spindle. Oocytes were analyzed using Imaris software (Oxford Instruments, England) and the National Institute of Health (NIH) ImageJ software (NIH, Bethesda, MD, USA).

### List of antibodies used for immunocytochemistry

Mouse anti-pericentrin (1:100, BD Biosciences, 611815), rabbit anti-pericentrin (1:100, Abcam, ab4448), rabbit anti-myosin X (1:100, Novus Biologicals, 22430002), rabbit anti-myosin VIIb (1:100, ProteinTech, 14467-1-AP), rabbit anti-myosin XVa (1:100, orb221541, Biorbyt Ltd), conjugated mouse anti-α-tubulin–Alexa Fluor 488 (1:100; Invitrogen, 322588), conjugated rabbit anti-α-tubulin–Alexa Fluor 488 (1:100; Invitrogen, 322588), anti-rabbit NuMA (1:100; Abcam, ab97585, rabbit anti-formin 2 (1:100; ProteinTech, 11259-1-AP) goat anti-mouse Alexa Fluor 350 (1:200; Invitrogen, A21049), goat anti-mouse Alexa Fluor 568 (1:200; Invitrogen, A10037), goat anti-mouse Alexa Fluor 647 (1:200; Invitrogen, A32728), goat anti-rabbit Alexa Fluor 568 (1:200; Invitrogen, A10042), and goat anti-rabbit Alexa Fluor 647 (1:200; Invitrogen, A32795)) and goat anti-rabbit Abberior Star 580 (1:100, Abberior GmbH, ST580-1002-500UG).

### Time-lapse confocal live imaging

Oocytes expressing fluorescently labeled proteins were imaged over time in milrinone-free CZB medium under a 40x, 63x or 100x oil immersion objective using Leica Stellaris confocal microscope equipped with a microenvironmental chamber (UNO-T-H-CO_2_, Okolab) to control temperature at 37°C in a humidified atmosphere containing 5% CO_2_. Image acquisition started as indicated in the corresponding figure legends. Live imaging settings were configured to span the entire oocyte or the region of interest at 0.5 to 5 μm Z-intervals depending on the specific parameter being measured. Smaller Z-steps were used to assess MyoX movement, F-actin growth and pMTOCs, whereas larger Z-steps were used to assess spindle migration, cytokinesis and PBE.

### In situ chromosome counting

*In vitro* matured oocytes were transferred to CZB medium containing 100 μM monastrol (MilliporeSigma, M8515) and incubated for an additional 2 hours. After monastrol treatment, the oocytes were fixed in a freshly prepared 2.5% paraformaldehyde solution and stained with CREST autoimmune serum (1:30, Antibodies Incorporated, 15-234) to label the kinetochores. DAPI was used to label the DNA. Fluorescence was observed using a 63× oil objective using Leica Stellaris confocal microscope, and oocytes were imaged using 0.5-μm Z-intervals. Oocytes were scored either as euploid (containing 40 kinetochores) or as aneuploid (containing greater or less than 40 kinetochores).

### Image processing and analysis

The number and volume of pMTOCs were processed and analyzed using the spot analysis and isosurface analysis features of Imaris software (Oxford Instruments). Briefly, according to^13^, based on pMTOC signal to noise fluorescence intensity, we adjusted the threshold value to the level at which all MTOCs are detected followed by MTOC surface segmentation. The numbers of pMTOCs were analyzed using the spot analysis feature, whereas pMTOC volume was analyzed using the isosurface analysis feature. The consistency of pMTOC number values obtained by the spot analysis was confirmed by calculating pMTOC number with the isosurface analysis. The same processing parameters were applied to each experimental analysis within the same replicate. The distribution of pMTOCs along the spindle axis was performed by a plot-profile graph using the ImageJ software.

To quantify cortical, cytoplasmic, and spindle-localized F-actin intensity, the laser power was adjusted to the level at which cytoplasmic F-actin was detected. To determine spindle-localized F-actin intensity or MyoX intensity, cytoplasmic F-actin/MyoX intensity was subtracted from spindle-localized F-actin/MyoX. To determine cytoplasmic and cortical F-actin intensity, multiple regions of interest (n=4) were quantified across the cytoplasm (not overlapping with the meiotic spindle) and the cortex, followed by background subtraction and, ultimately, averaged to denote a final F-actin intensity measurement for each oocyte. The same regions of interest area were fixed among all groups within the experiments. To quantify F-actin alignment, the longitudinal axis of the spindle was defined by drawing a line connecting the centers of the two spindle poles, which also passes through the center of the metaphase plate. Dispersion angle output was calculated using the “Directionality” plugin in ImageJ. A low dispersion indicates that the F-actin fibers are well aligned with the longitudinal axis of the spindle.

### Fluorescence recovery after photobleaching (FRAP)

FRAP was performed on oocyte expressing CEP192-mCherry and eGFP-EB3 or CEP192-mCherry and SiR-actin using Leica Stellaris confocal microscope equipped with a microenvironmental chamber under 100x oil immersion (1.4 NA). Bleaching was performed at pMTOC-associated MTs of metaphase I oocytes using 100% 488 nm laser power (100 ms laser pulse). After 10 seconds of imaging (2.6 frames per second), the region was photobleached and imaged for an additional 45 seconds (eGFP-EB3) or 5 min (SiR-actin). The photobleaching background was also measured in an area of equal size and adjacent to the photobleached region and was subtracted from the fluorescence intensity of the photobleached region in the spindle. The integrated FRAP Boaster option was enabled. The Automated fluorescence recovery curve was automatically generated using the integrated FRAP module in LAS X STELLARIS Control Software. Normalization was carried out by adjusting the data to the last prebleach time point. The data were corrected with a linear offset prior to the analysis for photobleaching.

### Quantitative RT-PCR

RNA from metaphase I mouse oocytes was isolated using the RNAqueous™ Micro Total RNA Isolation kit (Invitrogen) and cDNA was synthesized using the SuperScript™ IV Reverse Transcriptase (Invitrogen). Real-time PCR was performed with a QuantStudio 3 Real-Time PCR System (Applied BioSystems) using TaqMan® probes (ThermoFisher Scientific). Gapdh mRNA was used for normalization.

### List of TaqMan probes for qRT-PCR

MyoX (Mm00450859_m1), MyoVIIb (Mm00473514_m1), MyoXVa (Mm00465026_m1), and Gapdh (Mm99999915_g1).

### Western blot analysis

Metaphase I mouse oocytes (n=450) were washed in bicarbonate-free MEM followed by snap-freezing in liquid nitrogen. Oocytes were lysed in 1:1 dilution of MEM and 2X Laemmli Sample Buffer (Bio-Rad) supplemented with 1% β-mercaptoethanol and denatured at 95 °C for 10 min. The samples were separated by size in a 10% SDS polyacrylamide gel electrophoresis (Mini-PROTEAN TGX; Bio-Rad). A colorimetric ladder (PageRuler™ Plus Prestained Protein Ladder; Bio-Rad) was used. The proteins were then transferred to a 0.2μm nitrocellulose membrane using a Trans-Blot Turbo Transfer System (Bio-Rad). The membrane was incubated for 1 h in 2% blocking buffer [1X Tris-buffered Saline + 0.1% Tween 20 (TBS-T) + Blotting-Grade Blocker; Bio-Rad], followed by a 4 °C overnight incubation with primary antibodies. The membrane was washed 5 times with TBS-T and incubated for 1 h with a secondary antibody conjugated with horseradish peroxidase (KwikQuant Western Blot Detection Kit; Kindle BioSciences). After 5 washes with TBS-T, the ECL signal was detected with the Clarity Max™ Western ECL Substrate (Bio-Rad) in a ChemiDoc™ MP Imaging System (Bio-Rad).

### Statistical analysis

One-way analysis of variance (ANOVA) and Student t-test were used to evaluate the statistical differences between groups using the GraphPad Prism software. The one-way ANOVA test was followed by a Tukey post hoc test to allow the comparison and establish significant differences among groups. All experiments were repeated at least 3 times and the total number of analyzed oocytes per each group is specified above graphs. The differences of P < 0.05 were considered significant. The data is represented as means ± standard error of the mean (SEM).

### Data availability

No sequence or proteomic data has been generated in this study. All image data supporting the findings of this study are available from the corresponding author upon request. Source data are provided with this paper.

## Supplementary Figures

**Supplementary figure 1:**
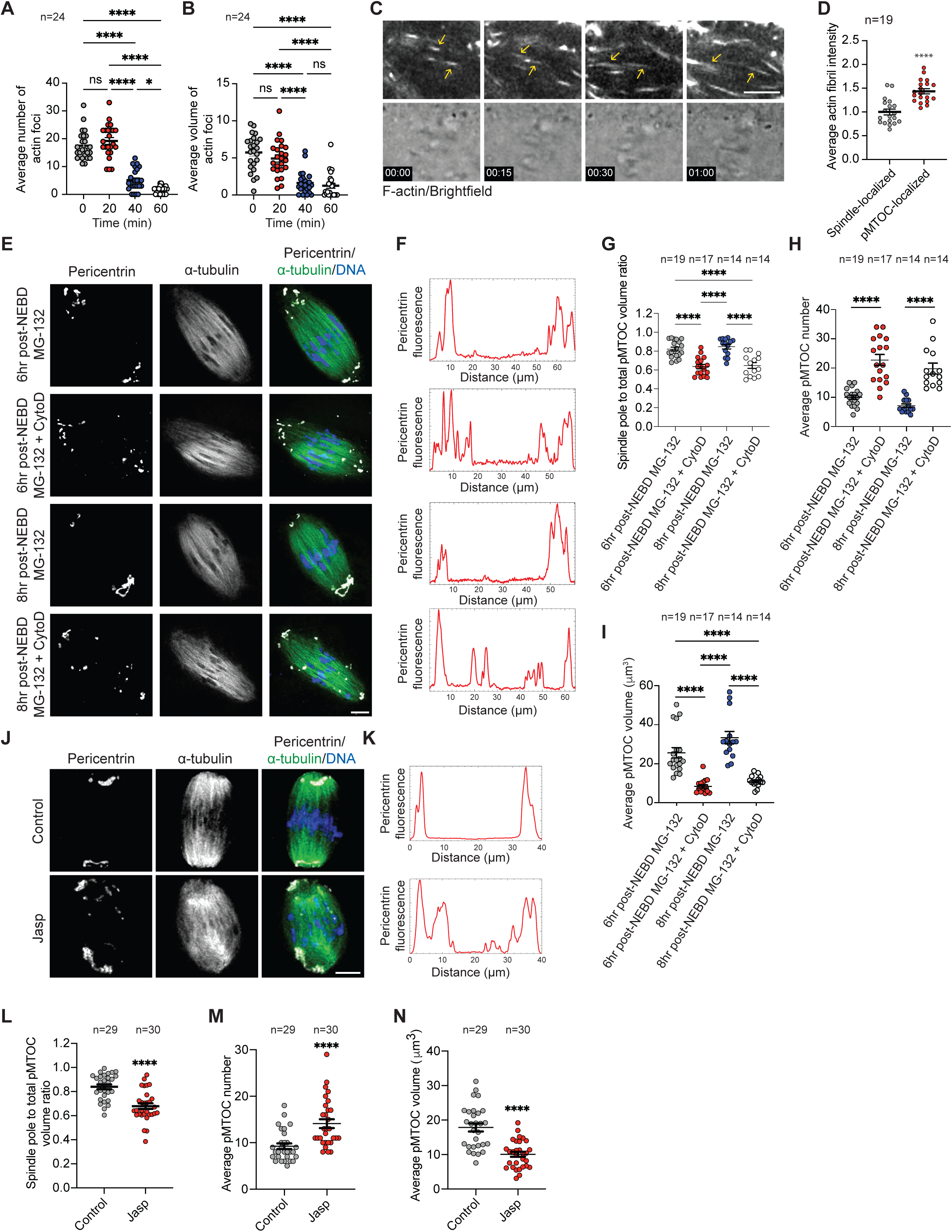
F-actin disruption leads to pMTOC sorting and clustering defect independent of a cell cycle delay. (**A**) Quantification of average number of actin foci after NEBD in Fig. 1D. (**B**) Quantification of average volume of actin foci after NEBD in Fig. 1D. (**C**) Representative images from time-lapse confocal microscopy of an oocyte stained with SiR-actin (F-actin, gray). Time points represent the timing from imaging. Yellow arrows represent F-actin fibril elongation. Scale bar represents 10 μm. The total number of examined oocytes from 3 independent replicates is 24 oocytes. (**D**) Quantification of average actin fibrils localized at pMTOCs or the spindle in metaphase I oocytes. (**E**) Representative confocal images of metaphase I oocytes treated with MG-132 (control), or MG-132 + CytoD at NEBD and fixed after 6 or 8 h post-NEBD. The oocytes were immunostained with α-tubulin to label the spindle (green) and pericentrin to label MTOCs (gray). DNA was labeled with 4′,6-diamidino-2-phenylindole, dihydrochloride (DAPI, blue). Scale bar represents 10 μm. (**F**) Plot profile of pMTOCs (pericentrin fluorescence) along the longitudinal axis of the spindle in the groups indicted in **E**. (**G**) Quantification of pMTOC volume at the spindle poles to the total volume of all pMTOCs in the spindle. (**H**) Average pMTOC number quantification after 3D reconstruction of pericentrin from oocytes in **E**. (**I**) Average pMTOC volume quantification from oocytes in **E**. (**J**) Representative confocal images of metaphase I oocytes treated with DMSO (control) or jasplakinolide (Jasp) at NEBD. Metaphase I oocytes were immunostained with α-tubulin to label the spindle (green) and pericentrin to label MTOCs (gray). DNA was labeled with 4′,6-diamidino-2-phenylindole, dihydrochloride (DAPI, blue). Scale bar represents 10 μm. (**J**) Plot profile of pMTOCs (pericentrin fluorescence) along the longitudinal axis of the spindle in the groups indicted in **J**. (**L**) Quantification of pMTOC volume at the spindle poles to the total volume of all pMTOCs in the spindle. (**M**) Average pMTOC number quantification in **J**. (**N**) Average pMTOC volume quantification after 3D reconstruction of pMTOCs from oocytes in **J**. Data are represented as means ± SEM. Asterisks represent significant differences, * p<0.05 and **** p<0.0001. The total number of analyzed oocytes per each experimental group (from 3 independent replicates) is specified above each graph.

**Supplementary figure 2:**
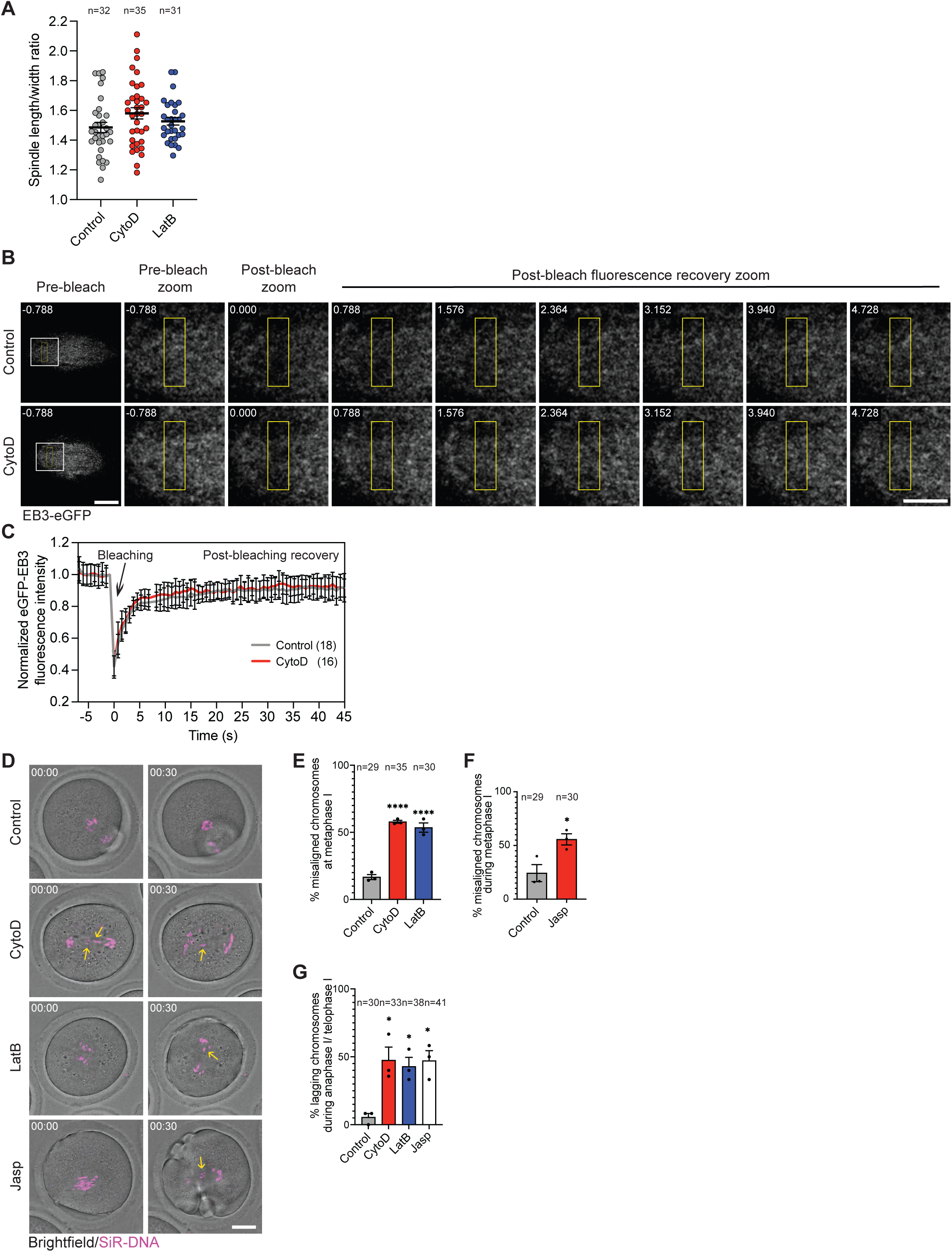
Disruption of F-actin leads to chromosome segregation errors. (**A**) Average spindle length to width ratio of oocytes treated with DMSO (control), Cytochalasin D (CytoD), or Latrunculin B (LatB). (**B**) FRAP was performed on oocyte expressing eGFP-EB3 and CEP192-mCherry and bleaching was performed at pMTOC-associated MTs of metaphase I oocytes using 100% 488 nm laser power (100 ms laser pulse) followed by imaging for an additional 45 seconds. Scale bar represents 10 μm. (**C**) Quantification of eGFP-EB3 fluorescence recovery after photobleaching (FRAP). (**D)** Time-lapse live confocal microscopy of mouse oocytes treated with DMSO (control), CytoD, LatB or jasplakinolide (Jasp) at NEBD and stained with SiR-DNA to label the DNA (pseudo magenta). Scale bar represents 20 μm. Chromosomes were tracked throughout live imaging by using the autotracking function of Imaris software. Yellow arrow heads represent lagging chromosomes. Time points represent the timing from the start of confocal imaging. (**E**) Quantification of average percentage of misaligned chromosomes at metaphase I in control, CytoD and LatB groups. (**F**) Quantification of average percentage of misaligned chromosomes at metaphase I in control and Jasp-treated oocytes. (**G**) Average percentage of lagging chromosomes at anaphase I/telophase I in the indicated groups. Data are represented as means ± SEM. Asterisks represent significant differences, * p<0.05 and **** p<0.0001. The total number of analyzed oocytes per each experimental group (from 3 independent replicates) is specified above each graph.

**Supplementary figure 3:**
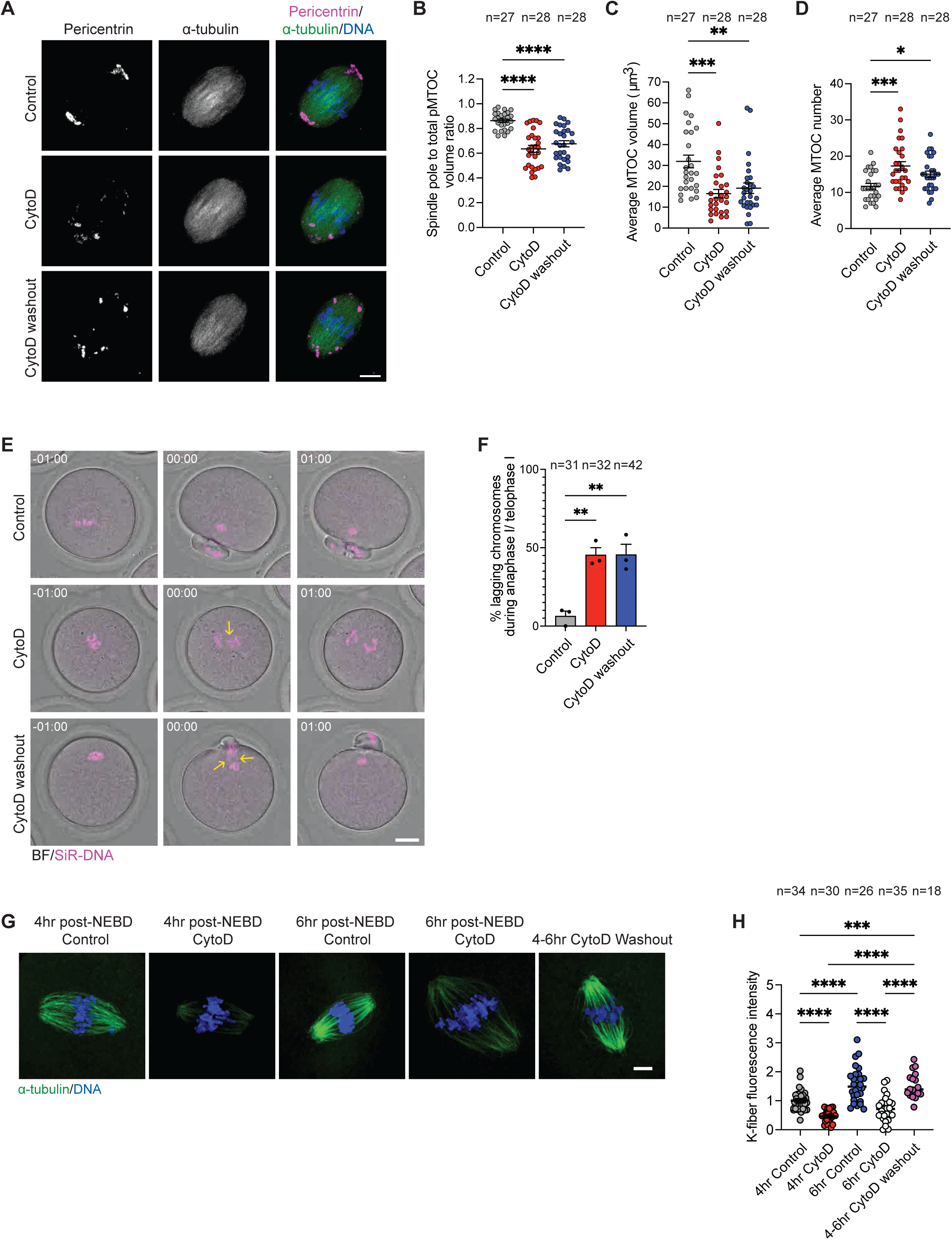
Acute disruption of F-actin at prometaphase I leads to chromosome missegregation. (**A**) Representative confocal images of metaphase I oocytes treated with DMSO (control), Cytochalasin D (CytoD) for 6 h or CytoD for 4 h followed by washing and culturing in CytoD-free medium for an additional 2 h (CytoD washout). Metaphase I oocytes (6 h-post NEBD) were immunostained with α-tubulin to label the spindle (green) and pericentrin to label MTOCs (magenta). DNA was labeled with 4′,6-diamidino-2-phenylindole, dihydrochloride (DAPI, blue). Scale bar represents 10 μm. (**B)** Quantification of pMTOC volume at the spindle poles to the total volume of all pMTOCs in the spindle. (**C**) Average pMTOC volume quantification after 3D reconstruction of pMTOCs from oocytes in the indicated groups. (**D)** Average pMTOC number quantification in the indicated groups. (**E)** Time-lapse live confocal microscopy of oocytes stained with SiR-DNA to label the DNA (pseudo magenta) from control, CytoD and CytoD washout groups at NEBD and stained with SiR-DNA to label the DNA (pseudo magenta) followed by time-lapse confocal microscopy. Scale bar represents 20 μm. Chromosomes were tracked throughout live imaging by using the autotracking function of Imaris software. Yellow arrow heads represent lagging chromosomes. Time points represent the timing from the start of confocal imaging. (**F**) Average percentage of lagging chromosomes at anaphase I/telophase I in the indicated groups. (**G**) Representative confocal images of control, CytoD and CytoD washout oocytes exposed to cold shock (10 min) at metaphase I (6 h-post NEBD) followed by immunostaining with α-tubulin to label the spindle (green). DNA was labeled with DAPI (blue). Scale bar represents 10 μm. (**H**) Average fluorescence intensity of kinetochore fibers (K-fibers). Data are represented as means ± SEM. Asterisks represent significant differences, * p<0.05, ** p<0.01, *** p<0.001 and **** p<0.0001. The total number of analyzed oocytes per each experimental group (from 3 independent replicates) is specified above each graph.

**Supplementary figure 4:**
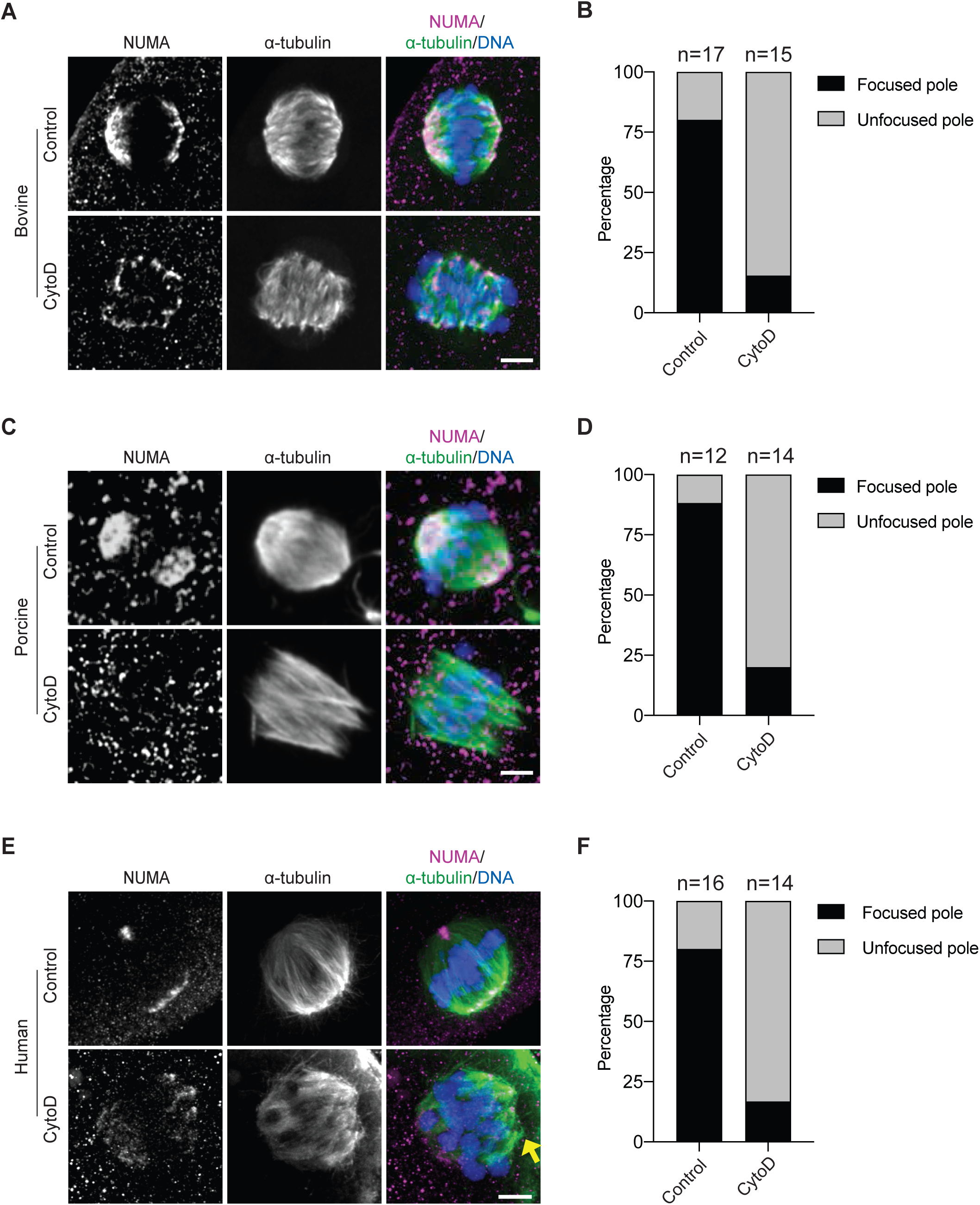
F-actin regulates spindle pole focusing in mammalian oocytes. Representative confocal images of metaphase I bovine **(A)**, porcine **(C)** and human **(E)** oocytes treated with DMSO (control) or CytoD at NEBD. Metaphase I oocytes were immunostained with α-tubulin to label the spindle (green) and NuMA (magenta). DNA was labeled with 4′,6- diamidino-2-phenylindole, dihydrochloride (DAPI, blue). Scale bar represents 10 μm. (**B, D, F**) Quantifications of spindle pole focusing in **A**, **C** and **E**, respectively. The total number of analyzed oocytes per each experimental group (from 3 independent replicates) is specified above each graph.

**Supplementary figure 5:**
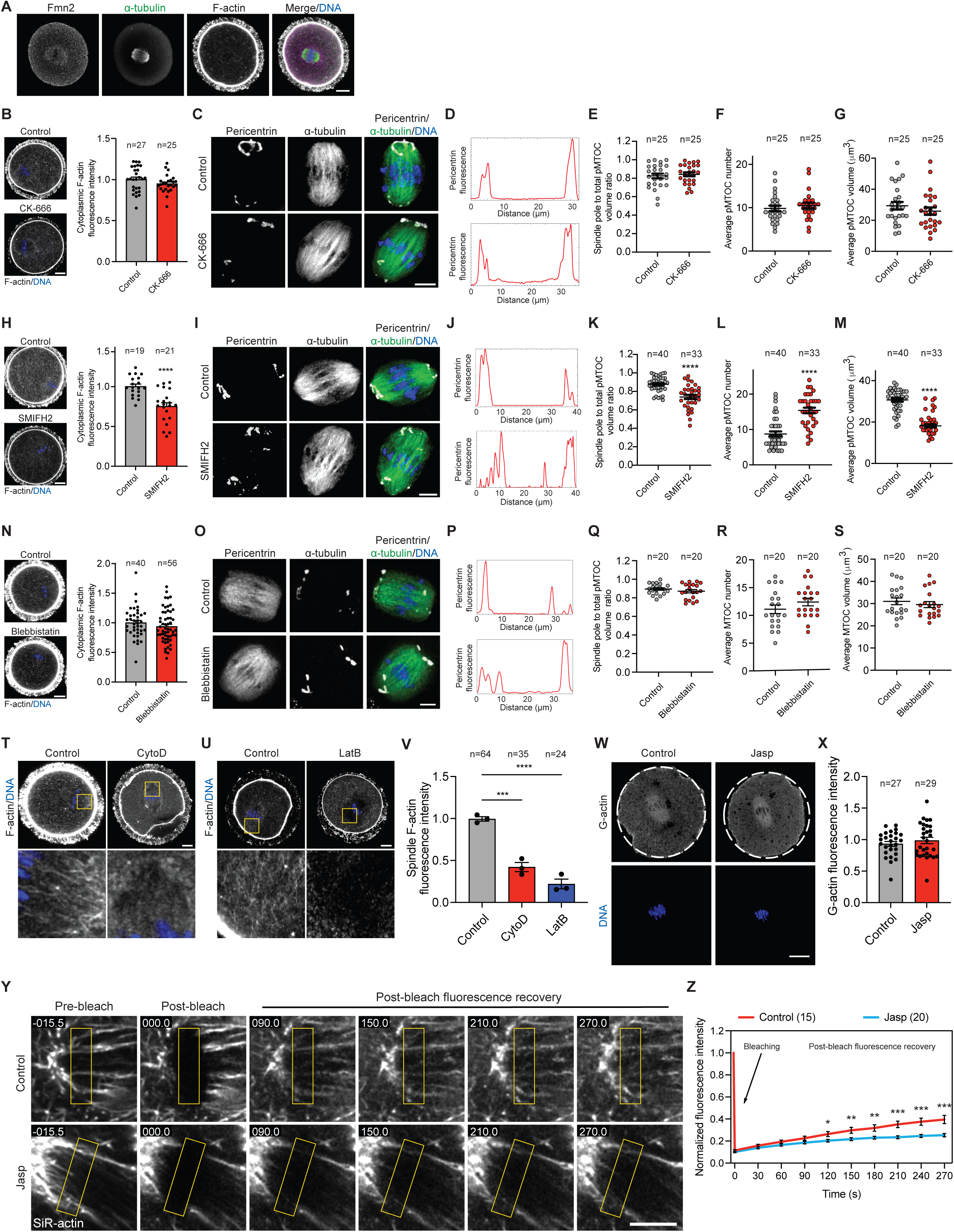
Disrupting cytoplasmic, but not cortical, F-actin results in pMTOC sorting and clustering defects. (**A**) Representative confocal images of a metaphase I oocyte immunostained with formin 2 (Fmn2, magenta) and α-tubulin (green) antibodies. DNA was labeled with 4′,6-diamidino-2- phenylindole, dihydrochloride (DAPI, blue) and F-actin was labeled with phalloidin (grey). Scale bar represents 20 μm. **(B,H,N)** Representative confocal images of oocytes stained with DAPI and phalloidin (left panels) and the corresponding quantification of cytoplasmic F-actin (right panels) in control, CK-666-, **B**), SMIFH2-, **H**) and Blebbistatin-, **N**) treated oocytes. **(C,I,O)** Representative confocal images of metaphase I oocytes treated with DMSO (control), CK-666 (**C**), SMIFH2 (**I**) or blebbistatin (**O**) at NEBD. Metaphase I oocytes were immunostained with α-tubulin to label the spindle (green) and pericentrin to label MTOCs (gray). DNA was labeled with DAPI (blue). Scale bar represents 10 μm. (**D,J,P)** Plot profile of pMTOCs (pericentrin fluorescence) along the longitudinal axis of the spindle in the indicated groups. (**E,K,Q)** Quantification of pMTOC volume at the spindle poles to the total volume of all pMTOCs at the spindle in the indicated groups. (**F,L,R)** Average pMTOC number quantification. (**G,M,S**) Average pMTOC volume quantification. (**T,U**) Representative confocal images of metaphase I oocytes treated at NEBD with DMSO (control), Cytochalasin D (CytoD), or Latrunculin B (LatB). The oocytes were stained with phalloidin (to label F-actin) and DAPI (to label the DNA). (**V**) Quantification of average spindle-localized F-actin. (**W**) Representative confocal images of metaphase I oocytes treated at NEBD with DMSO (control) or Jasplakinolide (Jasp). The oocytes were stained with DNase I antibody (to label G-actin). (**X**) Quantification of average G-actin in the indicated groups. (**Y**) Metaphase I oocytes treated with Jasp at NEBD were stained with SiR-actin to label F-actin. Bleaching was performed near the spindle poles using 100% 647 nm laser power (100 ms laser pulse) followed by imaging for an additional 5 minutes to monitor the fluorescence recovery after photobleaching (FRAP). (**Z**) Quantification of SiR-actin fluorescence recovery after photobleaching (FRAP). Data are represented as means ± SEM. Asterisks represent significant differences, * p<0.05, ** p<0.01, *** p<0.001 and **** p<0.0001. The total number of analyzed oocytes (from 3 independent replicates) is specified above each graph.

**Supplementary figure 6:**
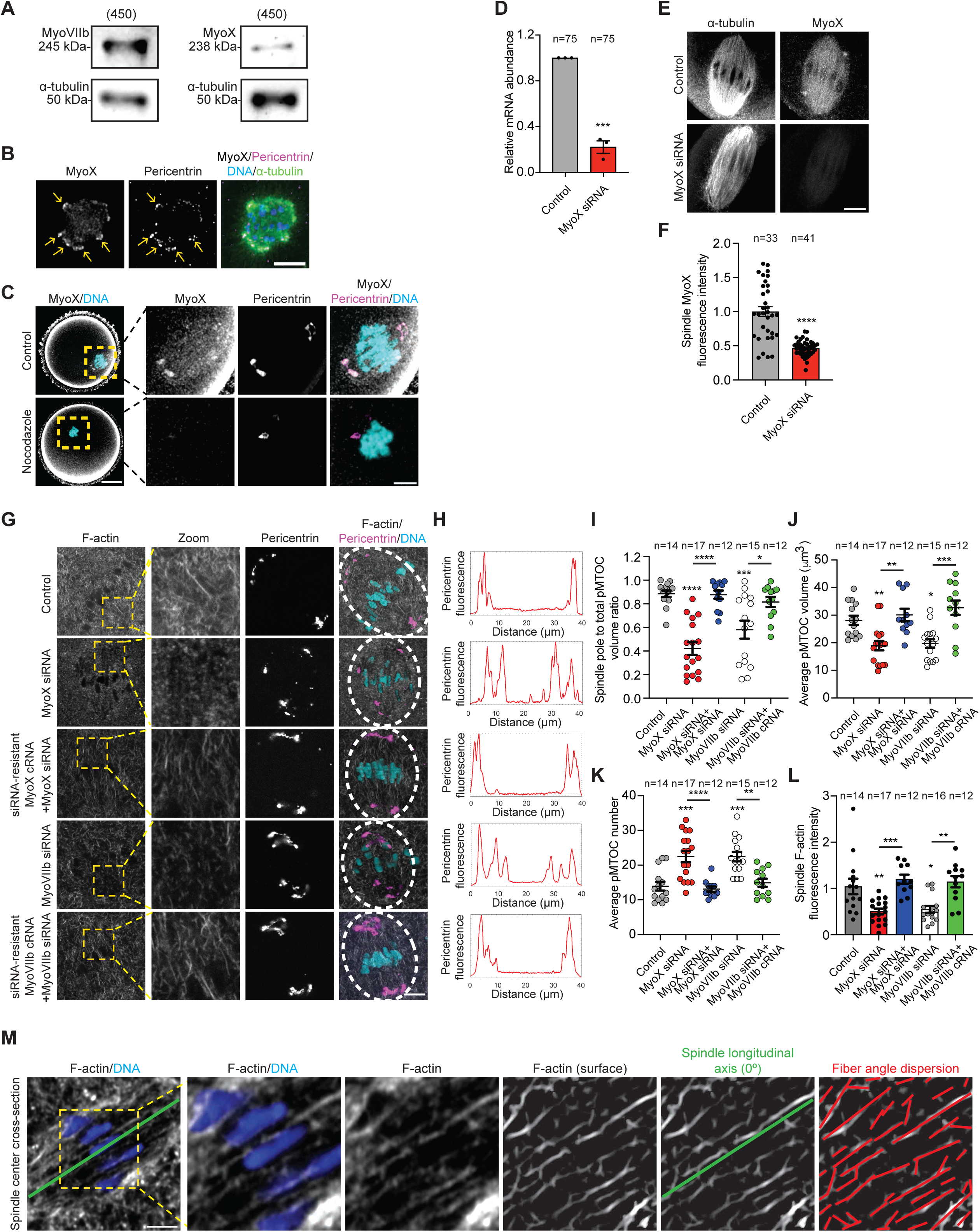
Myosin X localization to the spindle is microtubule dependent and its depletion perturbs pMTOC clustering and sorting. (**A**) Metaphase I oocytes (150 oocytes per group) were sampled and assessed by Western blot analysis with Myosin X and Myosin VIIb antibodies. (**B**) Representative confocal images of a prometaphase I oocyte immunostained with Myosin X (MyoX, gray), pericentrin (MTOCs, magenta) and α-tubulin (green) antibodies. DNA was labeled with 4′,6-diamidino-2-phenylindole, dihydrochloride (DAPI). Scale bar represents 10 μm. (**C**) Representative confocal images of metaphase I oocytes treated with DMSO (control) or nocodazole at NEBD. Metaphase I oocytes were immunostained with Myosin X (MyoX, gray) and pericentrin (MTOCs, magenta) antibodies. DNA was labeled with DAPI. Scale bar represents 10 μm. (**D**) 75 metaphase I oocytes from the indicated groups were isolated, pooled, and used for mRNA purification and generation of cDNA. Taqman probes were used to detect Myosin X mRNA transcript abundance. (**E**) Representative confocal images of metaphase I oocytes microinjected with control siRNA or Myosin X siRNA at the germinal vesicle stage. The oocytes were immunostained with α-tubulin and Myosin X antibodies. Scale bar represents 10 μm. (**F**) Quantification of average Myosin X fluorescence intensity at the spindle in **E**. (**G**) Representative confocal images of metaphase I oocytes from the indicated groups were immunostained with pericentrin (MTOCs, magenta) antibody and stained with phalloidin (F-actin) and DAPI (DNA). Scale bar represents 10 μm. (**H)** Plot profile of pMTOCs (pericentrin fluorescence) along the longitudinal axis of the spindle in the indicated groups. (**I)** Quantification of pMTOC volume at the spindle poles to the total volume of all pMTOCs at the spindle in the indicated groups. (**J**) Average pMTOC volume quantification after 3D reconstruction of pMTOCs from oocytes in the indicated groups. (**K)** Average pMTOC number quantification in the indicated groups. (**L**) Quantification of average spindle-localized F-actin in the indicated groups. (**M**) Representative images showing the method used to quantify F-actin alignment using the “Directionality” plugin in ImageJ. Data are represented as means ± SEM. Asterisks represent significant differences, * p<0.05, ** p<0.01, *** p<0.001 and **** p<0.0001. The total number of analyzed oocytes per each experimental group (from 3 independent replicates) is specified above each graph.

**Supplementary figure 7:**
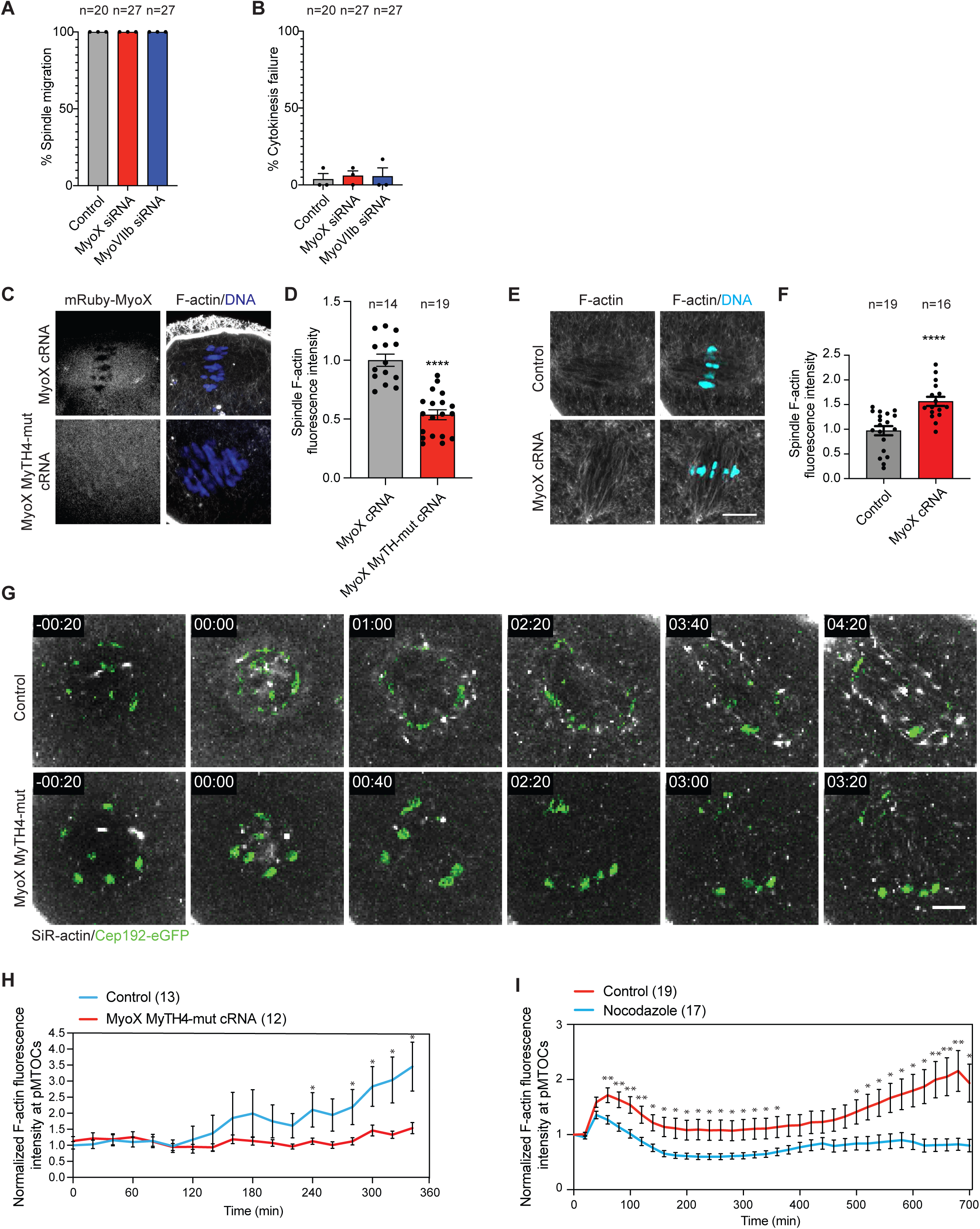
Myosin X recruits F-actin to the spindle through its microtubule binding domain. (**A,B)** Metaphase I oocytes microinjected with control siRNA, Myosin X siRNA or Myosin VIIb siRNA at the germinal vesicle stage were imaged live to assess the percentage of spindle migration (**A**) and the percentage of oocytes underwent cytokinesis failure (**B**). (**C**) Representative confocal images of metaphase I oocytes microinjected with mRuby-Myosin X full length (MyoX) or mRuby-Myosin X microtubule binding domain (MyTH4, K1651D/K1654D) mutant (MyoX MyTH-mut) at the germinal vesicle stage. The oocytes were stained with phalloidin (F-actin) and 4′,6-diamidino-2-phenylindole, dihydrochloride (DAPI, blue). Scale bar represents 10 μm. (**D**) Quantification of spindle F-actin fluorescence intensity in **C**. (**E**) Representative confocal images of metaphase I oocytes microinjected with mRuby or mRuby-Myosin X full length (MyoX) at the germinal vesicle stage. The oocytes were stained with phalloidin (F-actin) and DAPI (DNA, cyan). Scale bar represents 10 μm. (**F**) Quantification of spindle-localized F-actin fluorescence intensity in **E**. (**G**) Time-lapse live confocal microscopy of oocytes expressing Cep192-eGfp (pMTOCs, green) and stained with SiR-actin (F-actin, gray) and microinjected with mRuby or mRuby-Myosin X microtubule binding domain (MyTH4, K1651D/K1654D) mutant (MyoX MyTH-mut) at the germinal vesicle stage. Time points represent the timing from NEBD. Scale bar represents 10 μm. (**H**) Quantification of F-actin fluorescence intensity at polar MTOCs in **G**. (**I**) Quantification of F-actin fluorescence intensity at polar MTOCs in DMSO- and nocodazole-treated oocytes over time following NEBD. Data are represented as means ± SEM. Asterisks represent significant differences, * p<0.05, ** p<0.01 and **** p<0.0001. The total number of analyzed oocytes per each experimental group (from 3 independent replicates) is specified above each graph.

**Supplementary figure 8:**
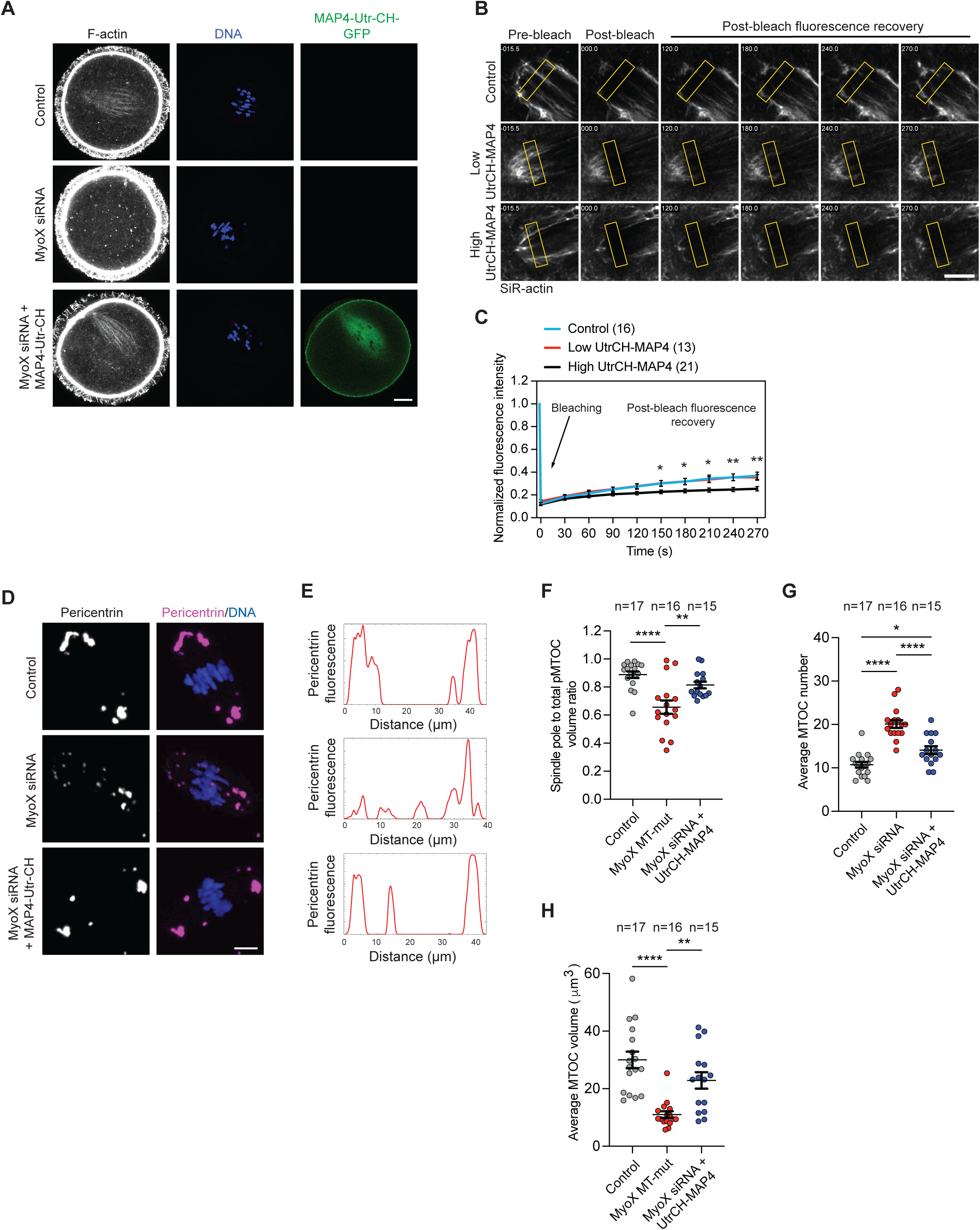
MAP4-UtrCH recruits F-actin to the spindle and rescues pMTOC clustering and sorting defects in Myosin X knockdown oocytes. (**A,D**) Representative confocal images of metaphase I oocytes microinjected with control siRNA, MyoX siRNA or MyoX siRNA + MAP4-UtrCH-GFP cRNA at the germinal vesicle stage. The oocytes were stained with phalloidin (F-actin). DNA was stained with 4′,6-diamidino-2-phenylindole, dihydrochloride (DAPI). Scale bar represents 20 μm. (**B**) Metaphase I oocytes, microinjected with 10 ng/μl (low concentration) or 100 ng/μl (high concentration) MAP4-UtrCH-GFP cRNA at the germinal vesicle stage, were stained with SiR-actin to label F-actin. Bleaching was performed near the spindle poles using 100% 647 nm laser power (100 ms laser pulse) followed by imaging for an additional 5 minutes to monitor the fluorescence recovery after photobleaching (FRAP). (**C**) Quantification of SiR-actin fluorescence recovery after photobleaching (FRAP). (**D**) Representative confocal images of metaphase I oocytes microinjected with control siRNA, MyoX siRNA or MyoX siRNA + MAP4-UtrCH-GFP cRNA at the germinal vesicle stage. The oocytes were immunostained with pericentrin antibody (pMTOCs). DNA was stained with DAPI. Scale bar represents 10 μm. (**E)** Plot profile of pMTOCs (pericentrin fluorescence) along the longitudinal axis of the spindle in the indicated groups. (**F**) Quantification of pMTOC volume at the spindle poles to the total volume of all pMTOCs at the spindle in the indicated groups. (**G)** Average pMTOC number quantification in the indicated groups. (**H**) Average pMTOC volume quantification after 3D reconstruction of pMTOCs from oocytes in the indicated groups. Data are represented as means ± SEM. Asterisks represent significant differences, * p<0.05, ** p<0.01 and **** p<0.0001. The total number of analyzed oocytes per each experimental group (from 3 independent replicates) is specified above each graph.

**Supplementary Figure 9:**
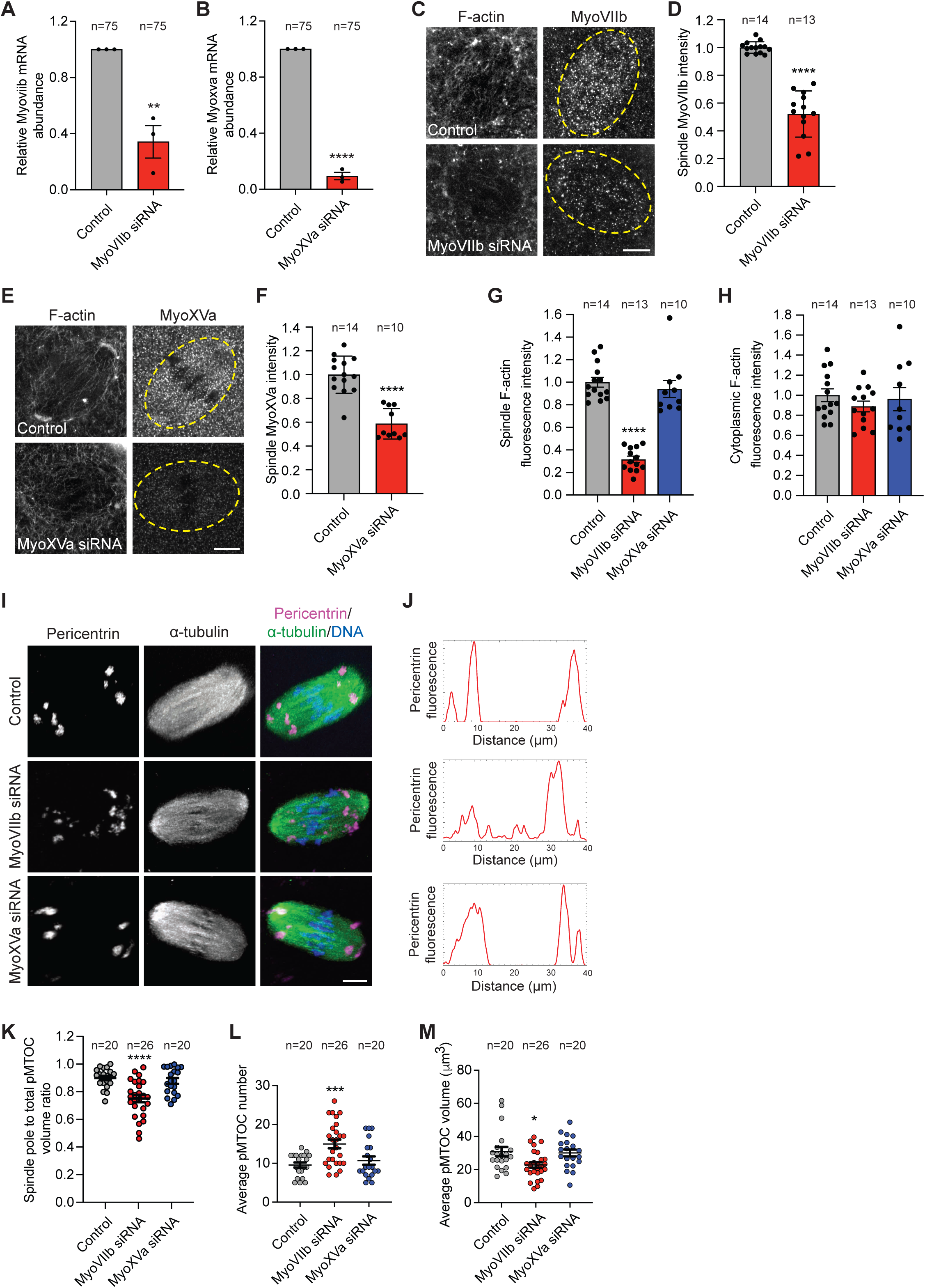
Myosin VIIb, but not XVa, regulates pMTOC sorting and clustering. (**A,B**) Sixty metaphase I oocytes from the indicated groups were isolated, pooled, and used for mRNA purification and generation of cDNA. Taqman probes were used to detect Myosin VIIb (**A**) or Myosin XVa (**B**) mRNA transcript abundance. (**C**) Representative confocal images of metaphase I oocytes microinjected with control siRNA or Myosin VIIb siRNA at the germinal vesicle stage. The oocytes were labeled with phalloidin (F-actin, gray) and Myosin VIIb antibody (gray). Scale bar represents 10 μm. (**D**) Average spindle-localized MyoVIIb fluorescence intensity in oocytes from **C**. (**E**) Representative confocal images of metaphase I oocytes microinjected with control siRNA or Myosin XVa siRNA at the germinal vesicle stage. The oocytes were labeled with phalloidin (F-actin, gray) and Myosin XVa antibody (MyoXVa, gray). Scale bar represents 10 μm. (**F**) Average spindle-localized MyoXVa fluorescence intensity in oocytes from **E**. (**G**) Average spindle-localized F-actin fluorescence intensity of metaphase I oocytes microinjected with control, MyoVIIb or MyoXVa siRNA at the germinal vesicle stage. (**H**) Average cytoplasmic F-actin fluorescence intensity of metaphase I oocytes microinjected with control, MyoVIIb or MyoXVa siRNA at the germinal vesicle stage. (**I**) Representative confocal images of metaphase I oocytes microinjected with control, MyoVIIb or MyoXVa siRNA at the germinal vesicle stage. Metaphase I oocytes were immunostained with α-tubulin to label the spindle (green) and pericentrin to label MTOCs (gray). DNA was labeled with 4′,6- diamidino-2-phenylindole, dihydrochloride (DAPI, blue). (**J**) Plot profile of pMTOCs (pericentrin fluorescence) along the longitudinal axis of the spindle in the groups indicted in **I**. (**K**) Quantification of pMTOC volume at the spindle poles to the total volume of all pMTOCs at the spindle in the groups indicated in **I**. (**L**) Average pMTOC number quantification after 3D reconstruction of pericentrin from oocytes in **I**. (**M**) Average pMTOC volume quantification after 3D reconstruction of pericentrin from oocytes in **I**. Data are represented as means ± SEM. Asterisks represent significant differences, * p<0.05, ** p<0.01, *** p<0.001 and **** p<0.0001. The total number of analyzed oocytes per each experimental group (from 3 independent replicates) is specified above each graph.

**Supplementary Figure 10:**
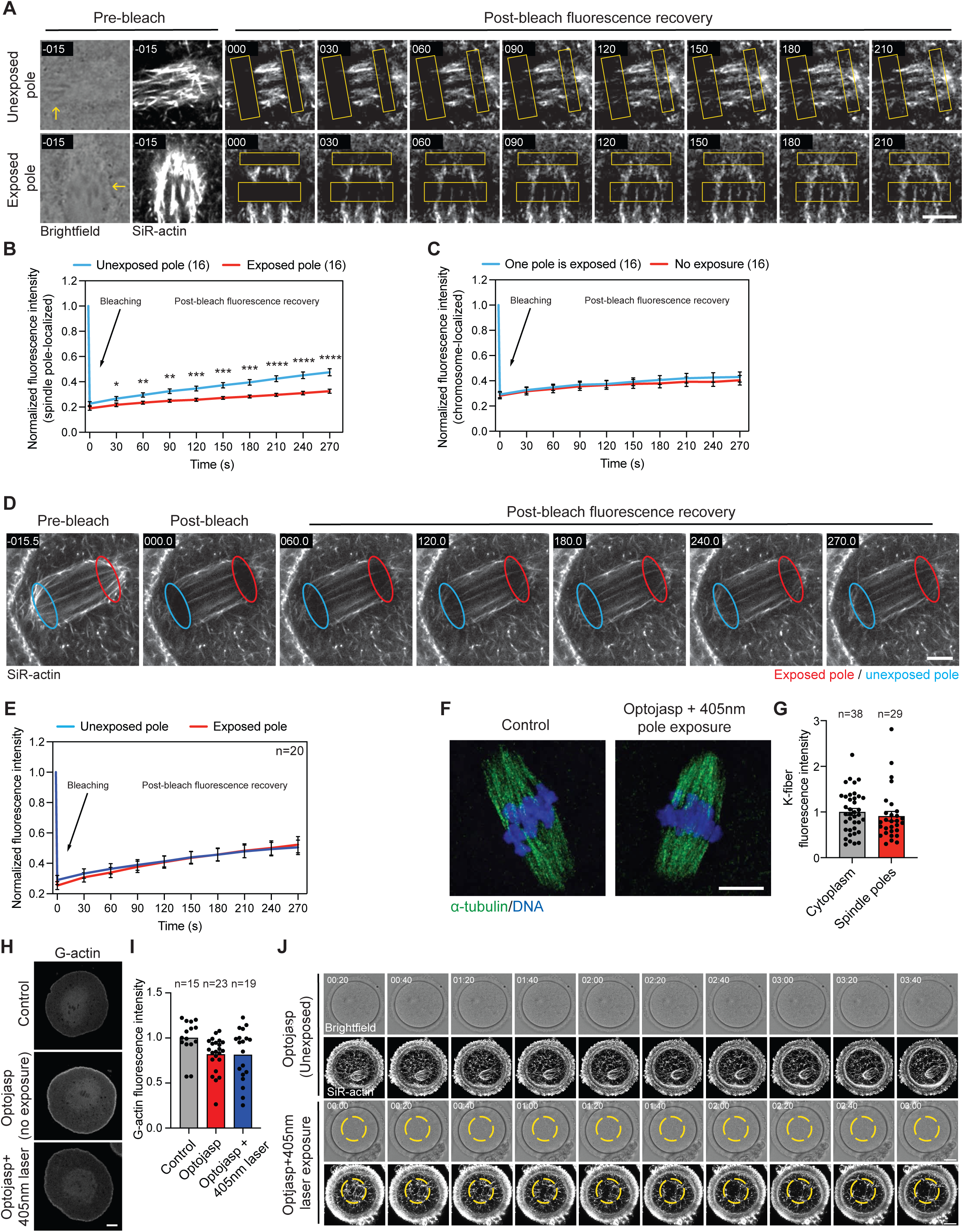
Optojasp-1 is an efficient tool to spatially disrupt F-actin in mouse oocytes. (**A**) Prometaphase I oocytes expressing CEP192-mCherry and labeled with SiR-actin (F-actin) were treated with Optojasp-1 and exposed to 405 nm laser at one spindle pole (exposed pole) or not exposed to 405 nm laser (no exposure). At metaphase I, bleaching of F-actin was performed at the exposed spindle pole, unexposed spindle pole or metaphase I chromosomes (in spindle pole-exposed oocytes or unexposed oocytes) using 100% 647 nm laser power (100 ms laser pulse) followed by imaging for an additional 5 minutes to monitor the fluorescence recovery after photobleaching (FRAP). (**B**) Quantification of SiR-actin fluorescence recovery after photobleaching (FRAP) at exposed and unexposed spindle poles. (**C**) Quantification of SiR-actin fluorescence recovery after photobleaching (FRAP) at chromosomes in exposed and unexposed oocytes. (**D**) Prometaphase I oocytes labeled with SiR-actin (F-actin) were treated with DMSO and exposed to 405 nm laser at one spindle pole (exposed pole). At metaphase I, bleaching of F-actin was performed at both spindle poles using 100% 647 nm laser power (100 ms laser pulse) followed by imaging for an additional 5 minutes to monitor the fluorescence recovery after photobleaching (FRAP). (**E**) Quantification of SiR-actin fluorescence recovery after photobleaching (FRAP) at exposed and unexposed spindle poles. (**F**) Optojasp-treated oocytes exposed to 405 nm laser at the cytoplasm (control) and optojasp-treated oocytes exposed to 405 nm laser at both spindle poles were examined for kinetochore-fiber (K-fiber) stability following cold shock (10 min). The oocytes were immunostained with α-tubulin to label the spindle (green). DNA was labeled with 4′,6-diamidino-2-phenylindole, dihydrochloride (DAPI, blue). Scale bar represents 10 μm. (**G**) Average fluorescence intensity of kinetochore fibers (K-fibers). (**H**) Representative confocal images of metaphase I oocytes exposed to the indicated treatments and stained with DNase I antibody (to label G-actin). DNA was labeled with DAPI. Scale bar represents 10 μm. (**I**) Average fluorescence intensity of G-actin in the indicated groups. (**J**) Time-lapse live imaging confocal microscopy of SiR-actin (F-actin, gray) in mouse oocytes treated with Optojasp-1 without 405 nm laser exposure (control) or with 405 nm laser exposure in a specific region of interest during prometaphase I. Images were taken every 20 min. Time points represent the timing from the start of confocal imaging. Scale bar represents 20 μm. Data are represented as means ± SEM. Asterisks represent significant differences, * p<0.05, ** p<0.01, *** p<0.001 and **** p<0.0001. The total number of analyzed oocytes per each experimental group (from 3 independent replicates) is specified above each graph.

**Supplementary Figure 11:**
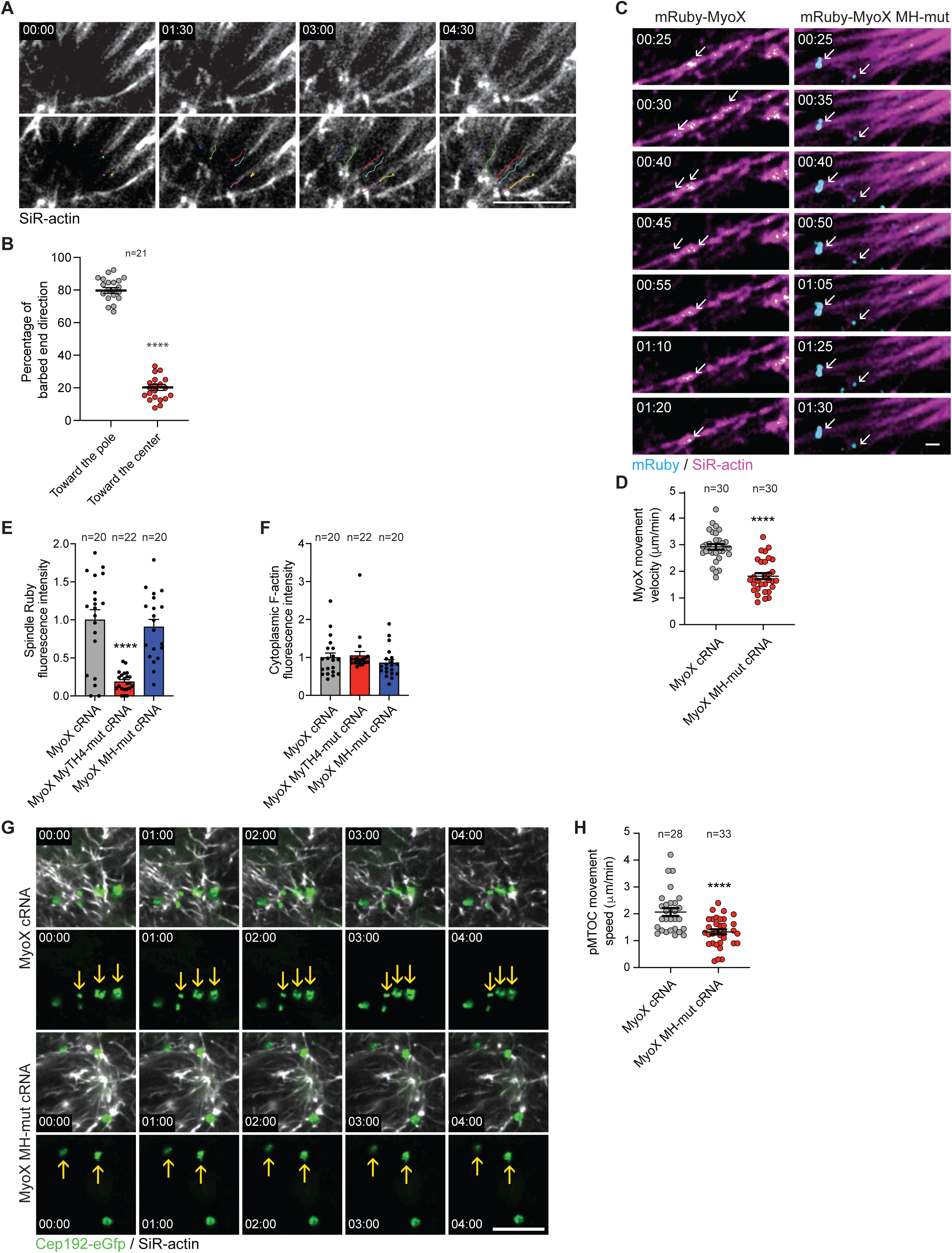
Myosin X lacking the head motor exhibits reduced motility on F-actin fibers and decreased pMTOC movement. (**A**) Metaphase I oocytes labeled with SiR-actin (F-actin) were exposed to photobleaching near the spindle pole using 100% 647 nm laser power (100 ms laser pulse) followed by imaging for an additional 5 minutes to monitor the direction of F-actin growth during the fluorescence recovery after photobleaching (FRAP). (**B**) Quantification of F-actin barbed end direction towards the spindle pole. (**C**) Time-lapse live imaging confocal microscopy of SiR-actin (F-actin, magenta) in mouse oocytes expressing mRuby fused to full-length mouse myosin X (mRuby-MyoX) or mRuby fused to myosin X lacking the head motor (mRuby-MH-mutant MyoX). Images were taken every 5 seconds. Time points represent the timing from the start of confocal imaging. White arrows represent the position of mRuby-MyoX and mRuby-MH-mutant MyoX over time. Scale bar represents 1 μm. (**D**) Quantification of Myosin X movement speed in **C**. (**E**) Quantification of average spindle-localized mRuby fluorescence intensity of oocytes microinjected with mRuby-tagged versions of MyoX, MyoX MyTH-mut, or MyoX MH-mut cRNA from Fig. 6B. (**F**) Quantification of average cytoplasmic F-actin fluorescence intensity of oocytes microinjected with mRuby-tagged versions of MyoX, MyoX MyTH-mut, or MyoX MH-mut cRNA from Fig. 6B. (**G**) Time-lapse live imaging confocal microscopy of SiR-actin (F-actin, gray) in Cep192-eGFP (MTOCs, green) mouse oocytes expressing myosin X (mRuby-MyoX) or mRuby fused to myosin X lacking the head motor (mRuby-MH-mutant MyoX). Images were taken every 5 seconds. Time points represent the timing from the start of confocal imaging. Yellow arrows represent the position of pMTOCs over time. Scale bar represents 10 μm. (**H**) Quantification of pMTOC movement speed in **G**. Data are represented as means ± SEM. Asterisks represent significant differences, **** p<0.0001. The total number of analyzed oocytes per each experimental group (from 3 independent replicates) is specified above each graph.

**Supplementary Figure 12:**
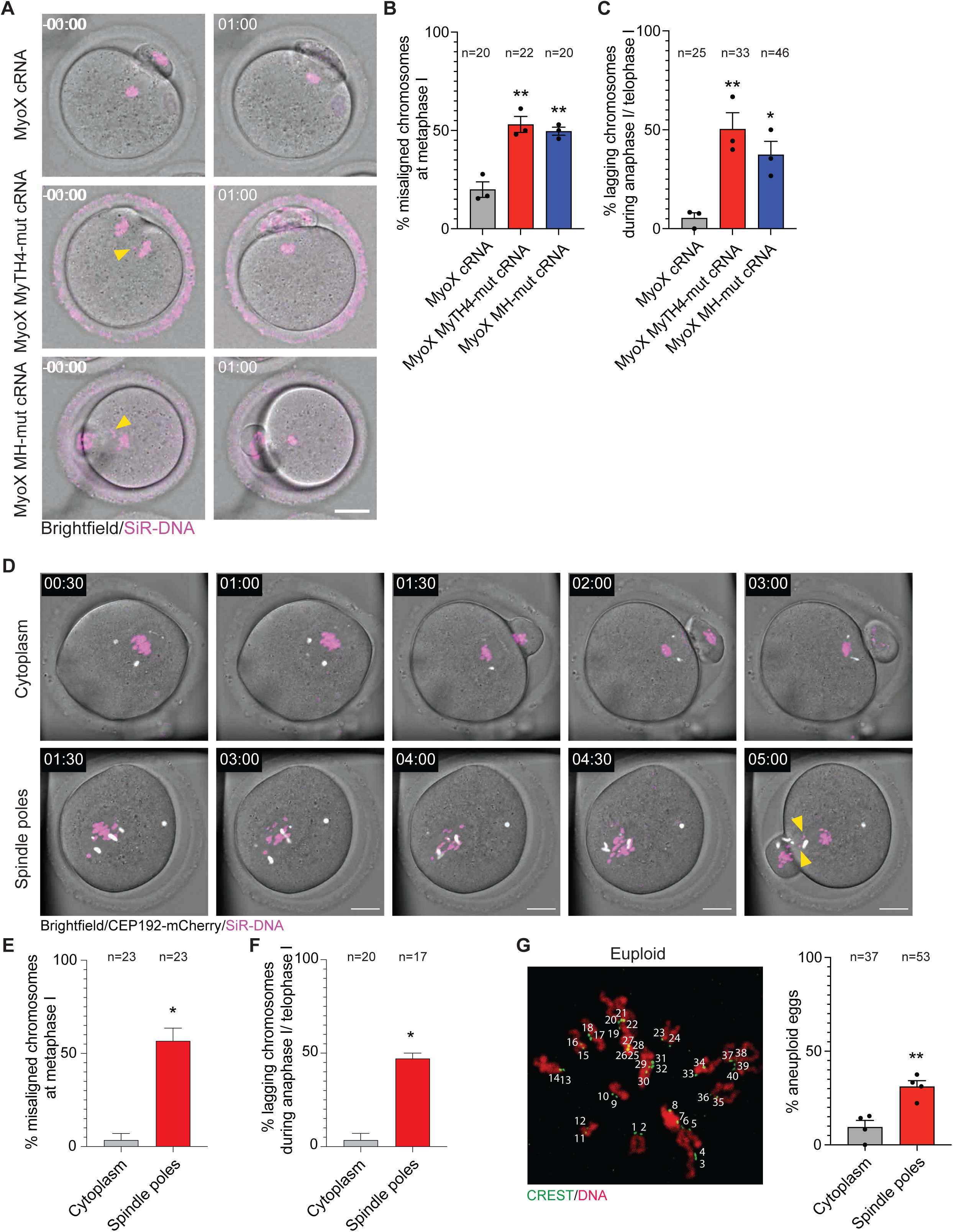
Disruption of spindle pole-localized cage-like F-actin leads to chromosome missegregation. (**A**) Time-lapse confocal microscopy of mouse oocytes expressing myosin X (MyoX), MyoX MyTH-mut, or MyoX MH-mut and stained with SiR-DNA to label the DNA. (**B**) Average percentage of misaligned chromosomes at metaphase I in the indicated groups. (**C**) Average percentage of lagging chromosomes at anaphase I/telophase I in the indicated groups. (**D**) Time-lapse confocal microscopy of MTOCs (Cep192-mCherry, pseudo gray) and DNA (SiR-DNA, pseudo magenta) in control and spindle-pole F-actin-perturbed oocytes. Prometaphase I mouse oocytes were treated with Optojasp-1 and exposed to 405 nm laser at the cytoplasm (control) or treated with Optojasp-1 and exposed to 405 nm laser at both spindle poles (spindle pole-localized F-actin perturbation). Images were taken every 30 min. Scale bar represents 20 μm. Chromosomes were tracked throughout live imaging by using the autotracking function of Imaris software. Yellow arrow heads represent lagging chromosomes. Time points represent the timing from the start of confocal imaging. (**E**) Average percentage of misaligned chromosomes at metaphase I of control and spindle pole F-actin-perturbed oocytes. (**F**) Average percentage of lagging chromosomes at anaphase I/telophase I of control and spindle pole F-actin-perturbed oocytes. (**G**) Representative images of euploid metaphase II oocytes assessed for aneuploidy by in situ chromosome counting. Quantification of the percentage of aneuploidy in the indicated groups (right panel). Data are represented as means ± SEM. Asterisk represents significant differences, * p<0.05, ** p<0.01, **** p<0.0001. The total number of analyzed oocytes per each experimental group from 3 independent replicates (except D-F, 2 replicates) is specified above each graph.

**Supplementary Movie 1: pMTOCs are associated with F-actin fibers.**

Time-lapse live confocal microscopy of pMTOCs (Cep192-eGfp, pseudo-magenta) and F-actin (SiR-actin, gray) in prometaphase I mouse oocytes. Fluorescence images were taken every 3 sec. Scale bar represents 5 μm.

**Supplementary Movie 2: Optojasp-1 is an efficient tool to spatially disrupt F-actin in mouse oocytes.**

Metaphase I oocytes expressing CEP192-mCherry (not imaged) and labeled with SiR-actin (F-actin) were treated with Optojasp-1 and exposed to 405 nm laser at one spindle pole (spindle pole-localized F-actin dynamic perturbation). Bleaching was performed 1 h post-laser exposure at the two spindle poles using 100% 647 nm laser power (100 ms laser pulse) followed by imaging for an additional 5 minutes to monitor the fluorescence recovery after photobleaching (FRAP). Scale bar represents 10 μm.

**Supplementary Movie 3: F-actin barbed ends are oriented towards the spindle poles.** Metaphase I oocytes labeled with SiR-actin (F-actin) were exposed photobleaching near the spindle pole using 100% 647 nm laser power (100 ms laser pulse) followed by imaging for an additional 5 minutes to monitor the direction of F-actin growth during the fluorescence recovery after photobleaching (FRAP). Scale bar represents 10 μm.

**Supplementary Movie 4: Polar MTOC clustering in oocytes expressing mRuby-Myosin X.** Time-lapse live imaging confocal microscopy of SiR-actin (F-actin, gray) in Cep192-eGFP (MTOCs, green) mouse oocytes expressing myosin X (mRuby-MyoX). Images were taken every 5 seconds. Time points represent the timing from the start of confocal imaging. Scale bar represents 10 μm.

**Supplementary Movie 5: Reduced polar MTOC movement in oocytes expressing mRuby-MH-mutant Myosin X.**

Time-lapse live imaging confocal microscopy of SiR-actin (F-actin, gray) in Cep192-eGFP (MTOCs, green) mouse oocytes expressing mRuby fused to myosin X lacking the head motor (mRuby-MH-mutant MyoX). Images were taken every 5 seconds. Time points represent the timing from the start of confocal imaging. Scale bar represents 10 μm.

## Acknowledgements

The authors would like to thank all members of the Glover lab and the Balboula lab for valuable help and discussions. The authors would like to thank Melina Schuh for providing the MAP4-UtrCH-GFP and Cep192-mCherry constructs. We thank Samantha Bosland, University of Missouri, for creating the schematic illustrations. We thank Alexander Jurkevich (the Molecular Cytology Core, University of Missouri) and Ivan Novotny (Light Microscopy Core Facility, Institute of Molecular Genetics of the Czech Academy of Sciences) for their support with the STED super-resolution imaging. This research was supported by the NIH (R35 GM142537) to AZB and Marie Sklodowska-Curie Fellowship 706170 (Horizon 2020, European Commission) to AZB and DMG. KK was supported by Czech Grant Agency (GA22-30494S). KK and JM were supported by Czech Health Research Council (NW24-08-00048), BIOCEV (CZ.1.05/1.1.00/02.0109), and Institute of Biotechnology of the Czech Academy of Sciences (RVO 86652036). HAD was funded by the Deutsche Forschungsgemeinschaft (DFG, German Research Foundation) under Germanýs Excellence Strategy – EXC 2051 – Project-ID 390713860. The Light Microscopy Core Facility, IMG, Prague, Czech Republic was supported by MEYS – LM2023050 and RVO – 68378050-KAV-NPUI.

## Author Contributions

D.M.G. and A.Z.B. conceived the project. E.S.-M., K.K., D.M.G. and A.Z.B. designed experiments. E.S.-M. and A.Z.B. analyzed experiments. E.S.-M. performed experiments. E.S.- M., N.N.A, M.F., O.T., J.M. performed human oocyte experiments. E.S.-M., N.N.A, M.F. performed STED oocyte imaging. V.N., F.K., D.T., H.-D.A., A.L.P., K.K., M.Z.-G., D.M.G. and A.Z.B. provided resources and discussed experiments. A.Z.B. supervised the work. E.S.-M. and A.Z.B. wrote the manuscript. All authors read and edited the manuscript.

